# Convergence of Independently Established Oncogenes in a Combined Breast and Ovarian Cancer Model: Hub Gene Validation and identification of potential Food-Derived therapeutics

**DOI:** 10.64898/2026.11.22.768457

**Authors:** Md. Jonaid Tawfiq, Md. Ismail Hossain Ashik, Chandan Barai, Ahmed Saif

**Author notes:** **Corresponding author:** Ahmed Saif, Department of Pharmacy, Faculty of Science, University of Rajshahi, Rajshahi 6205, Bangladesh.

## Abstract

Breast and ovarian cancers share molecular alterations that may reveal common therapeutic targets. Here, we integrated six Gene Expression Omnibus (GEO) datasets to identify shared transcriptional changes across the two cancers. Independent differential expression analysis followed by direction-specific intersection identified 396 shared differentially expressed genes (DEGs), comprising 192 upregulated and 204 downregulated genes. Protein-protein interaction analysis prioritized ten hub genes: AURKA, AURKB, BCL2, BIRC5, CCNA2, CDK1, FOXO1, KIF11, PTEN, and TOP2A. Functional enrichment highlighted cell-cycle regulation, mitotic spindle organization, and p53-associated signalling. Independent expression assessment using GEPIA2 identified seven hub genes significantly upregulated in both cancers, while FOXO1 was downregulated. Regulatory network analysis further identified candidate transcription factors and miRNAs potentially associated with these hubs. For therapeutic exploration, a 70,478-compound FooDB library was progressively filtered to 233 candidates using physicochemical, structural-alert, machine-learning-based bioactivity, and ADMET criteria. Molecular docking against eight selected druggable hubs, followed by decoy benchmarking and 100-ns molecular dynamics simulations, prioritized three complexes: CDK1-artonin L, AURKB-taraxacin, and KIF11-L-thyronine. CDK1-artonin L demonstrated the strongest overall computational support, whereas the AURKB and KIF11 complexes showed differing levels of binding and energetic evidence.

This study provides an integrated multi-cohort framework for identifying shared molecular hubs in breast and ovarian cancer and prioritizing food-derived compounds for further investigation. The findings nominate CDK1, AURKB, and KIF11 for experimental evaluation.

## 1. Introduction

Breast and ovarian cancer are two of the most common and most lethal cancers in women. Breast cancer is by far the larger burden: in 2022 it caused about 2.3 million new cases and 666,000 deaths worldwide, making it the most frequently diagnosed cancer in women and the leading cause of their cancer deaths in more than a hundred countries.^1,2^ Ovarian cancer is rarer but proportionally more dangerous, with roughly 324,000 diagnoses and 207,000 deaths in the same year.^1,3^ The two also behave differently at presentation. Breast cancer incidence climbs from the mid-forties and the disease is often caught early through screening;^2^ ovarian cancer is usually found only after it has spread, because no reliable screening test exists for women at average risk.^3^

The two diseases begin at different ages, but once underway they behave alike in ways that matter clinically: both recur frequently, both acquire resistance to chemotherapy, both metastasise, and both become hard to cure once advanced.^4,5^ Treatment has improved without solving this. Surgery, platinum–taxane chemotherapy, endocrine therapy and PARP inhibition all control the disease better than before, yet none removes the recurrence that defines both.^4,5^ In some patients the two cancers even share a genetic origin: inherited BRCA1 or BRCA2 mutations sharply raise the lifetime risk of either, which is the basis of hereditary breast–ovarian cancer syndrome.^6^ The connection runs deeper than inheritance. The two share a measurable genetic architecture, with common risk variants concentrated in cell-cycle and cell-death genes,^7^ and their most aggressive forms resemble one another transcriptionally, with high-grade serous ovarian tumors closely mirroring basal-like breast cancer.^8^

This overlap is not limited to one pathway. It runs through several shared programs, some now shown in direct comparisons rather than only inferred from separate studies. In both cancers the disease often starts the same way, with an early TP53 mutation that inactivates the p53 pathway.^9^ Once p53 is lost, the CDK–cyclin machinery that controls entry into mitosis is no longer restrained, and the cell-cycle hub genes flagged in breast-cancer studies keep reappearing in independent ovarian-cancer studies.^10,11^ The mitotic spindle and chromosome-segregation machinery fail alongside it, producing the mis-segregation, chromosomal instability and aneuploidy seen in both; spindle genes such as the Aurora kinases and kinesin motors are picked out as hubs in breast and ovarian cohorts alike.^12–14^ Genome maintenance is the next shared weakness: defective homologous-recombination repair, usually through BRCA1/2, underlies the hereditary syndrome and leaves both cancers sensitive to PARP inhibition.

Survival signalling is shared as well. The PI3K–AKT–mTOR axis is altered in both diseases: PIK3CA mutation and PTEN loss are common in breast cancer,^15^ and PI3K-pathway changes appear in a substantial fraction of ovarian tumors.^16^ Active AKT phosphorylates the tumor suppressor FOXO1 and forces it out of the nucleus, shutting down its control of cell-cycle arrest and apoptosis;^17^ loss of PTEN removes the main brake on the same pathway. Both FOXO1 and PTEN tend to be down-regulated in these cancers. Apoptosis is held off in part by the anti-apoptotic proteins survivin (BIRC5) and BCL2,^18^ and survivin turns up as a prognostic hub gene in ovarian^19^ and breast^20^ cancer alike. Wnt/β-catenin and Notch signalling add a further shared layer, driving epithelial–mesenchymal transition, cancer-stem-cell maintenance and chemoresistance: Wnt activity tracks with grade, EMT and platinum resistance in ovarian cancer,^21^ and with proliferation, metastasis and stemness in breast cancer.^22^

None of these genes is new to either cancer on its own, and we do not claim to have discovered them; the open question is how the shared signal has been studied. Past work falls into roughly three groups. Most of it is single-disease screens that, run separately, keep landing on the same mitotic and cell-cycle hubs. CDK1, CCNA2, TOP2A, KIF11 and the Aurora kinases recur across independent breast cohorts^10,11^ and, separately, across independent ovarian cohorts.^23,24^ The agreement is suggestive but only inferred: the shared signal is read off two parallel gene lists rather than drawn from one pooled analysis, so it cannot tell a real cross-tumor dependency from coincidence. A smaller body of work integrates the two diseases directly, but each only partway. One stays within the tumor stroma,^25^ and others fold breast and ovarian into panels of several tumor types,^26^ where averaging across cancers blurs whatever is specific to this pair. Others apply the same approach to a neighboring pairing instead, such as endometriosis–ovarian cancer.^27^

Two gaps follow from this. First, no one has isolated the whole-tumor signal shared specifically by breast and ovarian cancer in a single pipeline; it is always inferred across separate papers, confined to one compartment or one layer of biology, or diluted across unrelated tumors. Second, even the studies that integrate the two properly tend to stop at listing regulators or querying drug-repurposing databases,^26^ without taking the targets through to structure-based screening. That leaves two basic questions open: which of these shared hubs hold up as reproducible, externally validated signals rather than artifacts of a single dataset, and how any of them might actually be drugged. Answering these questions calls for an integrated workflow that analyses multiple GEO datasets across both diseases, identifies consistently dysregulated genes through within-cancer and cross-cancer intersection analyses, builds a protein–protein interaction network to prioritize hub genes, reconstructs their upstream regulatory networks, validates their expression in independent patient cohorts, and advances the prioritized targets towards structure-based drug screening.

A ranked list of hub genes still does not tell us what controls them. Because hub genes are driven by shared transcription factors and tuned by microRNAs,^28^ rebuilding the transcription-factor–gene–miRNA network is what exposes the upstream nodes coordinating the shared program.

These same genes also help tumors survive treatment. In triple-negative breast cancer, chemotherapy itself drives Aurora-kinase levels up and makes the tumor resistant to paclitaxel, which is what puts AURKB forward as a marker for stratifying patients.^14^ In ovarian cancer, inhibiting AURKA lowers PARP and BRCA1/2, and targeting AURKA and PARP together is synergistic in both BRCA-mutant and BRCA-wild-type models.^29^ Survivin keeps cells alive by blocking the apoptosis that chemotherapy is meant to trigger,^18^ while the FOXO–FOXM1 switch shapes the response to taxanes, anthracyclines and PI3K/AKT inhibitors.^30^

Targeting the over-expressed hubs does not have to start from nothing. Natural products are a large, chemically varied pool of candidate inhibitors with a long record in cancer-drug discovery, and they suit computational screening well;^31^ food-derived compounds add the further advantages of accessibility and tolerability. Testing thousands of them against several targets in the laboratory is impractical, so we used machine-learning models trained on bioactivity data to rank compounds and predict potency (IC50) before any docking, narrowing the library to a workable shortlist for docking and molecular-dynamics evaluation.^32^ We used this step cautiously. IC50 models lose accuracy on chemically unfamiliar compounds,^33^ and food-derived scaffolds are exactly that, so the predictions served to prioritize compounds for docking rather than as final estimates of binding affinity.

In this study we pool several GEO cohorts of breast and ovarian carcinoma to identify the genes both dysregulate, build a protein–protein interaction network from them, and prioritize ten hub genes — AURKA, AURKB, BCL2, BIRC5, CCNA2, CDK1, FOXO1, KIF11, PTEN and TOP2A — that cover the mitotic, DNA-repair, apoptotic and PI3K–AKT–FOXO programs above. We reconstruct the transcription-factor and miRNA networks behind these hubs and test whether their expression and prognostic value hold up in independent TCGA and GEPIA data. The over-expressed, druggable hubs are then screened against food-derived chemicals from FooDB, with machine-learning bioactivity prediction as a pre-docking filter ahead of molecular-dynamics confirmation; FOXO1 and PTEN we keep as regulatory and prognostic markers rather than drug targets. The point is the integration, not the individual genes: a reproducible, multi-cohort, cross-cancer pipeline that brings together shared transcriptomic analysis, regulatory-network reconstruction, external prognostic validation and machine-learning-guided natural-compound prioritization. The overall workflow is summarized in Figure 1.

**Figure 1.**
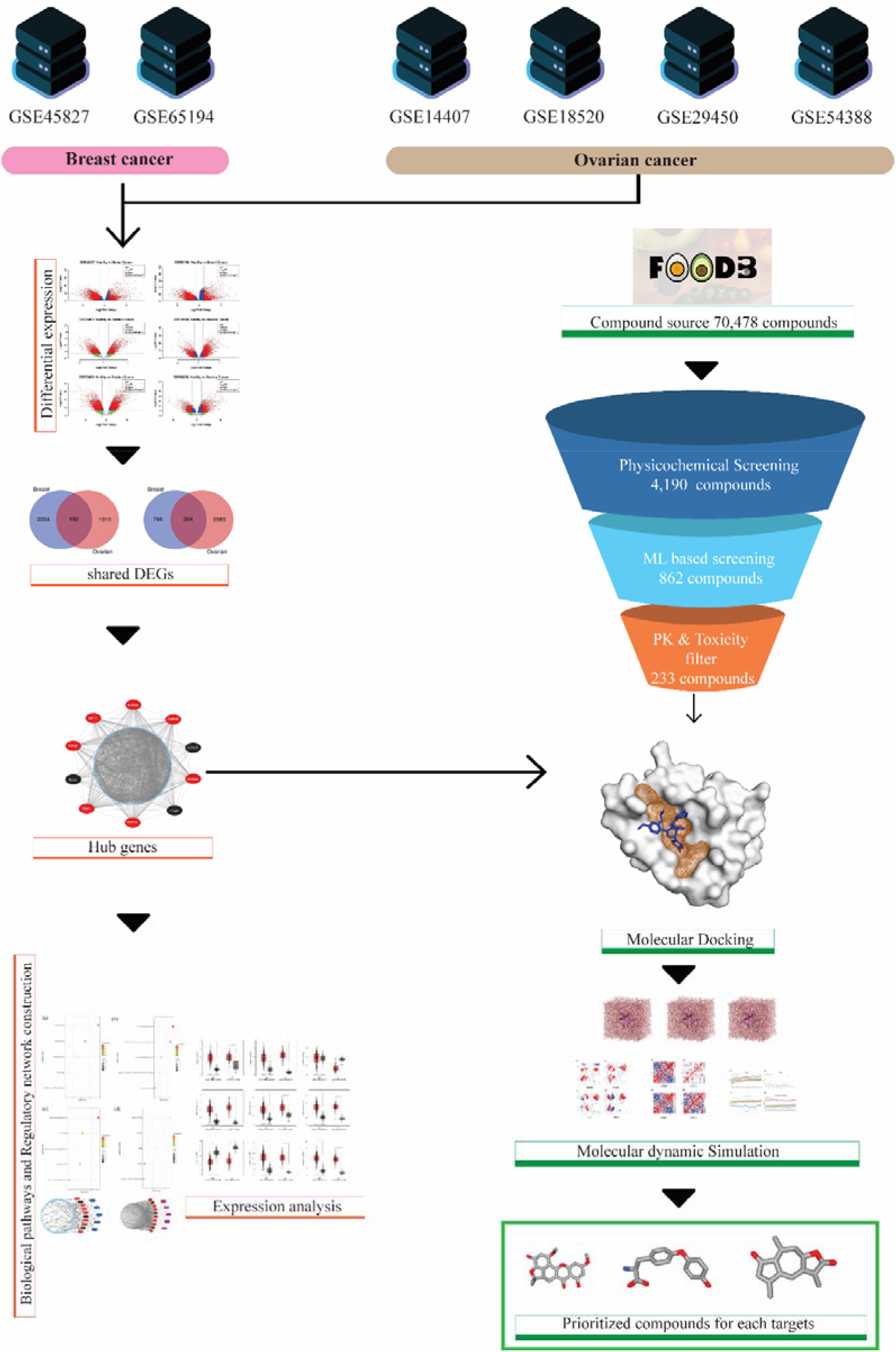
Integrated bioinformatics and drug discovery workflow identifying shared hub genes in breast and ovarian cancer, followed by compound screening, molecular docking, and molecular dynamics simulations to prioritize potential target-specific drug candidates.

## 2. Materials and Methods

### 2.1. Dataset acquisition and preprocessing

This study is based on the analysis of six transcriptomic microarray datasets retrieved from the Gene Expression Omnibus (GEO) repository^34^. Specifically, datasets GSE45827^35^ and GSE65194^36–38^ were used to examine breast cancer, and datasets GSE14407^39^, GSE18520^40^, GSE29450, and GSE54388^41^ were employed for the investigation of ovarian cancer. Investigation of transcriptomic profiles led to identification of genomic biomarkers and target receptors that can serve as potential therapeutic candidates.

### 2.2. Identification of differentially expressed genes (DEGs)

Identifying viable molecular targets for new drug interventions hinges on the analysis of Differentially Expressed Genes (DEGs). These genes exhibit statistically significant changes in expression under different conditions, and characterizing their dysregulated profiles is essential for understanding the mechanisms of a disease.^42^ For differential expression analysis, we utilized the “limma” package in R, a popular and robust tool for microarray and high-throughput data. A key strength of Limma is its use of linear models, which accommodates complex experimental designs and improves statistical power, even with small sample sizes. The package provides a comprehensive workflow, including tools for data normalization and exploration, and also allows for higher-level gene set analysis to better interpret the biological significance of expression changes.^43^ Limma employs a moderated t-statistic, where the standard errors are shrunk towards a common value, to calculate p-values that assess the significance of differential expression between case and control groups

These p-values were subsequently adjusted for multiple comparisons using the Benjamini-Hochberg method to control the false discovery rate.^44^ A gene was identified as a DEG based on its adjusted p-value and its average log2 fold-change (aLog2FC) between the two conditions. The criteria were defined as follows:

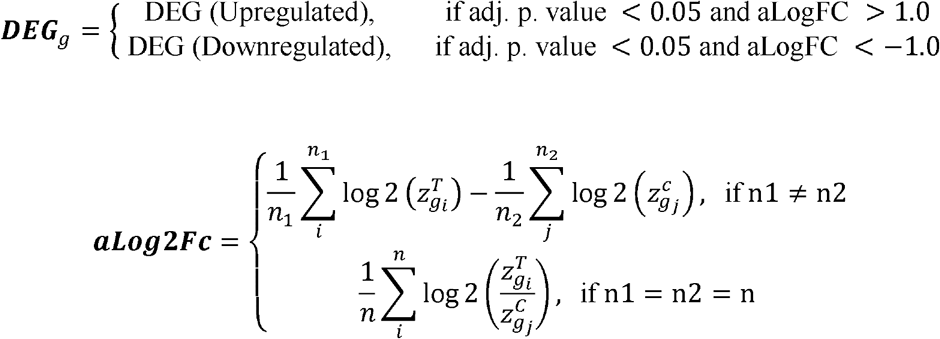

Here,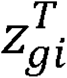and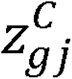represent the expression levels for the g_th_ gene in the i_th_ treatment and j_th_ control samples, respectively. We then identified upregulated and downregulated DEGs separately for each of the six datasets, and subsequently determined the common upregulated and downregulated genes, which were subsequently merged into a final set of common differentially expressed genes (cDEGs) for each disease.

### 2.3. Identification of shared DEGs (sDEGs)

We performed separate intersections for upregulated and downregulated genes to ensure that the identified cDEGs shared not only statistical significance but also the same direction of dysregulation, strengthening their biological relevance to a common pathogenic mechanism. To identify genes with a consistent pattern of dysregulation across both breast and ovarian cancer, a multi-step intersection analysis was performed. First, a robust set of DEGs was established for each disease individually. For breast cancer, we identified the DEGs that were common across all its datasets. An analogous procedure was performed for ovarian cancer to find the DEGs consistently identified across all of its respective datasets. This initial step was conducted separately for upregulated and downregulated genes to preserve the direction of regulation.

Subsequently, the resulting disease-specific DEG lists were compared to identify the final set of common DEGs (cDEGs). The list of common upregulated DEGs from breast cancer was intersected with the common upregulated list from ovarian cancer. A parallel intersection was performed for the downregulated gene lists. The genes identified in these final intersections were termed as shared differentially expressed genes (sDEGs) representing genes that are robustly and commonly dysregulated in both malignancies.

### 2.4. Identification of shared key genes (sKGs)

To translate our list of common DEGs into a set of key functional players (sKGs), we first modeled their relationships within a protein-protein interaction (PPI)^45^ network sourced from the STRING-v11 database.^46^ For identifying the most central nodes in this network, we turned to the cytoHubba^47^ plugin for Cytoscape^48^. This tool allowed us to apply a multi-faceted approach, assessing each gene’s importance through five different topological metrics: Degree^49^, Betweenness^50^, BottleNeck^51^, Closeness^52^ and Stress^53^. The distance ***D*** between two proteins (***u*** and ***v***) was calculated based on their respective neighbor sets, ***Nu*** and ***Nv***.

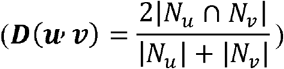

### 2.5. Functional enrichment and pathway analysis

The identified gene list was subjected to functional enrichment analysis using the DAVID database (https://david.ncifcrf.gov/), with ‘OFFICIAL_GENE_SYMBOL’ selected as the identifier and *Homo sapiens* specified as the background organism. KEGG pathway enrichment was performed using the Functional Annotation Chart module.

Statistical significance was determined using a modified Fisher’s Exact Test (EASE score), with a threshold set at *P* < 0.05. KEGG pathways were specifically selected due to their well-curated representation of cancer-related signaling networks. Finally, significant pathways were visualized using the visualizer platform (https://www.bioinformatics.com.cn/en) to facilitate comprehensive interpretation of functional patterns.

### 2.6. Regulatory network analysis of sKGs

To elucidate the transcriptional and post-transcriptional regulatory landscape of the selected key genes (sKGs), we constructed a gene regulatory network (GRN) utilizing the NetworkAnalyst 3.0 platform ^54^.

For transcriptional regulation, transcription factor (TF)-gene interactions were retrieved from the ChEA3 database (ChIP-X Enrichment Analysis 3)^55^, which integrates ChIP-seq data to ensure high-confidence binding evidence. For post-transcriptional regulation, miRNA-gene interactions were mapped using experimentally validated data from TarBase v9.0.^56^

To identify the most influential regulators (Hubs), the resulting network topology was analyzed based on Degree Centrality and Betweenness Centrality. Nodes with the highest connectivity (degree ≥ 7 for transcription factors and ≥ 8 for miRNAs) were identified as key regulatory drivers. The final TF-miRNA-mRNA triple-regulatory network was visualized using Cytoscape v3.9.1.^48^

### 2.7. Gene expression analysis

To validate the expression levels of the sKGs, the Gene Expression Profiling Interactive Analysis (GEPIA2) platform (http://gepia2.cancer-pku.cn/) was used to examine transcriptomic data from The Cancer Genome Atlas (TCGA). Expression levels of the ten sKGs (AURKA, AURKB, KIF11, BIRC5, CCNA2, CDK1, TOP2A, FOXO1, BCL2, and PTEN) were compared between tumor and matched normal tissues in breast invasive carcinoma (BRCA; tumor, n = 1085; normal, n = 291) and ovarian serous cystadenocarcinoma (OV; tumor, n = 426; normal, n = 88). Expression values were presented as log2(TPM + 1), and statistical significance between tumor and normal groups was determined using a one-way ANOVA, with a disease state considered a variable for calculating differential expression.

### 2.8. Collection of potential lead compounds

Food-derived compounds have emerged as promising lead candidates in the context of pharmacotherapy. Their appeal stems from several converging advantages. First, these compounds demonstrate multitarget bioactivities capable of simultaneously disrupting multiple oncogenic pathways, making them well suited to address the complexity of cancer biology.^57^ Second, their extensive history of human dietary consumption provides substantial preclinical safety data, reducing early-stage toxicity concerns compared to entirely novel synthetic scaffolds.^58^ Third, decades of epidemiological research have consistently linked plant-rich dietary patterns to 20-40% reductions in breast/ovarian cancer risk, providing robust clinical correlations.^59–61^ Finally, their isolation from abundant dietary sources via established extraction protocols substantially lowers discovery and scale-up costs compared to de novo synthesis.

The primary compound library underpinning this work is FooDB.^62^ This resource catalogs more than 80,000 unique food constituents from various food sources.

From a practical standpoint, most of the phytochemicals of interest can be obtained through straightforward extraction procedures from commonly consumed foods including cruciferous vegetables, berries and culinary spices. Established synthetic routes further support scale-up to pharmaceutical quantities without exceptional manufacturing demands. Among the most instructive examples is sulforaphane, derived principally from broccoli sprouts. Clinically, it has demonstrated the capacity to reverse cisplatin resistance in ovarian cancer xenograft models, a mechanism linked to upregulation of miR-30a-3p and concurrent downregulation of the DNA repair genes ERCC1 and ATP7A.^63^ Separately, sulforaphane has been shown to suppress metastatic behavior in triple-negative breast cancer, specifically in MDA-MB-231 xenografts, by attenuating signalling through the RAF/MEK/ERK pathway.^64^

Taken together, these lines of evidence position the systematic screening of dietary phytochemicals as an approach that is simultaneously scientifically rigorous and practically viable for advancing drug discovery efforts in breast and ovarian cancer.

### 2.9. Ligand library preparation

A large database of potential lead compounds were retrieved from the FooDB database.^62^ The downloaded structures were standardized and subjected to a multi-tiered computational filtration pipeline using RDKit^65^. Initially, compounds were screened against established drug-likeness and physicochemical property filters, including Lipinski’s Rule of Five^66^, Ghose^67^, Veber^68^, Egan^68^, and Muegge^69^ filters. Compounds flagged by PAINS (Pan-Assay Interference Compounds)^70^, Brenk^71^ structural alerts, or NIH chemical filters were excluded to eliminate potentially promiscuous or reactive scaffolds. Compounds passing all filters were subsequently evaluated for ADMET (Absorption, Distribution, Metabolism, Excretion, and Toxicity) properties using Deep-PK^72^, a deep-learning platform that employs graph neural networks to predict pharmacokinetic and toxicity endpoints. Specifically, predictions for AMES mutagenicity, carcinogenicity, and drug-induced liver injury (DILI) were assessed, and only compounds classified as safe across all three endpoints were retained for downstream studies.

### 2.10. Machine learning-based prediction of compound bioactivity

To reduce the filtered phytochemical library to a number tractable for molecular docking, a ligand-based machine-learning model was trained on experimental inhibition data and applied as a pre-docking prioritization filter.

Quantitative bioactivity data were retrieved from the ChEMBL database (release 37)^73^for the five hub targets with sufficient experimental data to support model training: AURKA (CHEMBL4722), AURKB (CHEMBL2185), CDK1 (CHEMBL308), KIF11 (CHEMBL4581) and TOP2A (CHEMBL1806). The three remaining targets subsequently carried into docking were not represented in the training set, as CCNA2 (CHEMBL2582), BIRC5 (CHEMBL5989) and FOXO1 (CHEMBL5294) returned only 8, 7 and 21 IC50 measurements with a defined pChEMBL value, respectively, which is too few to support model development.

Compound structures and their experimentally reported bioactivity measurements were retrieved and processed prior to machine-learning analysis. Initially, an individual machine-learning model was developed separately for each protein. For each target, the available ligand–bioactivity dataset was processed and divided into training and testing datasets. The same machine-learning workflow was applied consistently across all targets. The trained models were then evaluated using the coefficient of determination (R²) and root mean squared error (RMSE). To investigate whether information obtained from several biologically related targets could improve predictive performance, a second strategy was developed. Instead of training the model independently for each protein, the curated compound–bioactivity datasets from CDK1, AURKB, KIF11, TOP2A anf AURKA were combined to generate a larger pooled dataset. The combined dataset was subsequently used to train a single machine-learning model using the same general modeling and validation strategy. The trained model was then applied to the FooDB library, and compounds with a predicted pIC50 above 5, corresponding to a predicted IC50 below 10 µM, were retained for ADMET screening and subsequent docking.

### 2.11. Molecular docking

Protein-ligand docking was employed to investigate the binding mechanisms, preferred orientations, and interaction profiles of the selected phytochemical compounds against eight target proteins: AURKA (PDB: 1MQ4)^74^, AURKB (PDB: 4AF3)^75^, BIRC5 (PDB: 1XOX)^76^, CCNA2 (PDB: 4BCN)^77^, CDK1 (PDB: 4Y72)^78^, FOXO1 (PDB: 8A62)^79^, KIF11 (PDB: 3K5E), and TOP2A (PDB: 9BQ6)^80^, retrieved from the RCSB Protein Data Bank^81^. Prior to docking, protein structures were prepared by removing water molecules, heteroatoms, and co-crystallised ligands, followed by energy minimization using the GROMOS96^82^ force field implemented in Swiss-PDB Viewer (v4.1.0)^83^. Ligand structures obtained in SDF format were subjected to geometry using the MMFF94 force field^84^ and protonated at physiological pH 7.4 to reflect biologically relevant ionisation states. Structures were subsequently converted to PDBQT format using Open Babel. Molecular docking was performed using AutoDock Vina (v1.2.0)^85^. The top-ranked binding pose with the lowest predicted binding free energy (ΔG, kcal/mol) was selected for each ligand-protein pair. Binding interactions, including hydrogen bonds, hydrophobic contacts, and electrostatic interactions, were visualized and analysed using BIOVIA Discovery Studio Visualizer.

### 2.12. Docking validation

A decoy-based validation was performed to test whether the docking protocol could correctly distinguish true binders from non-binders. For each of the three prioritized ligands (CID 5241825, CID 44258662 and CID 5461103), 50 property-matched decoy molecules were generated by using a custom Python script based on the LUDe approach. These decoys were selected from the ChEMBL database using specific criteria to match the active ligands: Molecular Weight (MW) ± 20 Da, LogP ± 0.5, Hydrogen Bond Acceptors (HBA) ± 2, Hydrogen Bond Donors (HBD) ± 1 and number of Rotatable Bonds (RB) ± 2. Finally, to make sure the decoys were structurally different from the active ligands, any molecule with a Tanimoto coefficient of 0.30 or higher was removed.

This decoy-generation procedure follows the LIDeB Useful Decoys (LUDe) strategy^86^, in which property-matched but topologically dissimilar molecules serve as presumed non-binders. Decoy construction and the dissimilarity threshold followed the published LUDe specification^86^. Each decoy set was docked against its corresponding receptor using grid dimensions and scoring parameters identical to those applied to the active ligands. Docking performance was then expressed, for each target, as the Z-score of the reference ligand relative to the mean and standard deviation of its decoy score distribution, together with the ligand’s percentile rank and the number of decoys achieving a more favorable score.

#### 2.12.1. Redocking of co-crystallised ligands

To test whether the docking procedure reproduces experimentally observed binding modes, the co-crystallised ligand of each target was redocked into its own receptor: LZ9 into CDK1 (PDB 4Y72), VX6 into AURKB (PDB 4AF3) and K5E into KIF11 (PDB 3K5E) using the ame docking protocol.

#### 2.12.2. Benchmarking against known inhibitors

To test whether docking scores separate compounds with measured activity from compounds without, and to place the three prioritised ligands on that scale, a benchmark set was assembled for CDK1 (CHEMBL308), AURKB (CHEMBL2185) and KIF11 (CHEMBL4581) from ChEMBL release 37.^73^ IC50 records with a defined pChEMBL value and an exact (“=”) relation were retrieved, and replicate measurements of the same parent structure (first 14 characters of the InChIKey) were averaged. Compounds with a molecular weight outside 200–650 Da or with more than 50 heavy atoms were excluded. Known actives were defined as pChEMBL ≥ 7 (IC50 ≤ 100 nM) and measured inactives as pChEMBL ≤ 5 (IC50 ≥ 10 µM). Forty compounds per class and target were drawn at random (seed 0) before any docking was performed, giving 80 benchmark compounds per target. The largest fragment of each structure was converted to a three-dimensional conformer and minimised with MMFF94 in RDKit, prepared with Meeko with all rings held rigid, and docked into the same receptor, site-centred box and Vina version as in Section 2.12.1 (exhaustiveness 16, fixed seed). The three prioritised ligands were docked identically into all three receptors. Discrimination was quantified as the area under the receiver operating characteristic curve (ROC-AUC) using the negated Vina score as predictor, with a 95% confidence interval from 2,000 bootstrap resamples.^87^ Because docking scores tend to increase with ligand size,^88^ the AUC obtained from heavy-atom count alone was calculated as a control. The position of each prioritised ligand was expressed as the percentage of known actives and of measured inactives of the same target that it out-scored.

### 2.13. Molecular dynamics simulation

The functional activity of proteins is intrinsically linked to their capacity for significant conformational dynamics, which encompass rigid body movements, localized structural rearrangements, allosteric regulation, and global fold switching.^89–91^ Molecular dynamics (MD) simulations are employed to investigate the dynamic behavior of proteins, such as their flexibility and stability, by modeling the conformational changes that occur under conditions mimicking a physiological environment, including variable pH, salt concentrations, and the explicit presence of water and ions.

In this study, molecular dynamics (MD) simulations were performed using the GROMACS 2024.1^92^ The protein topology was generated using the CHARMM36 all-atom force field (charmm36-jul2022.ff)^93^ whereas the ligand topologies were constructed via the CGenFF server.^94^ Each system was explicitly solvated in a periodic boundary box with the TIP3P^95^ water model and neutralized with counter-ions. Prior to conducting molecular dynamics simulations, energy minimization was performed on each system using the steepest descent algorithm for up to 50,000 steps until the maximum force fell below 1000 kJ mol^−1^ nm^−1^. Following minimization, each protein-ligand complex was subjected to a two-step equilibration protocol for 100 ps in both canonical (NVT) and isothermal-isobaric (NPT) ensembles under position restraints on heavy atoms. The system temperature was maintained at 300K using the velocity-rescaling (V-rescale) thermostat^96^ and a pressure of bar was maintained during NPT equilibration using the Berendsen barostat^97^. Following equilibration, unrestrained 100 ns production runs were conducted. Trajectories were subsequently analyzed using standard GROMACS utilities.

### 2.14. Binding free-energy calculation (MM/GBSA)

Binding free energies of the CDK1–artonin L, AURKB–taraxacin and KIF11–L-thyronine complexes were estimated by the molecular mechanics/generalised Born surface area (MM/GBSA) method with gmx_MMPBSA v1.6.5.^98^ The single-trajectory approach was applied, in which receptor and ligand energies are taken from the complex trajectory. Two hundred frames, evenly spaced (every 50th frame of the analysed trajectory), were evaluated at 298.15 K. Polar solvation was calculated with the generalised Born model and the non-polar contribution from LCPO surface areas.^99^ The binding free energy was computed as ΔG_bind_ = ΔE_vdW_ + ΔE_ele_ + ΔG_GB_ + ΔG_SA_ = ΔG_gas_ + ΔG_solv_. Conformational entropy was not included, so the values are used to rank complexes and are not interpreted as absolute affinities.^100^ Values are given as the mean ± standard deviation (SD) over frames. Because consecutive frames are correlated, the lag-1 autocorrelation of ΔG_bind_ was used to estimate an effective number of independent frames (n_eff_ = n(1 − ρ)/(1 + ρ)) and the corresponding standard error of the mean (SEM), and the means of the first and second halves of the frames were compared to assess drift. Per-frame energies were taken from the gmx_MMPBSA output files; the component sums were checked (ΔG_gas_ + ΔG_solv_ = ΔG_bind_) before use.

## 3. Results

### 3.1. Identification of differentially expressed genes (DEGs)

For breast cancer, the process yielded 2,416 upregulated and 1000 downregulated genes (Figure 2). The analysis of ovarian cancer revealed a distinct profile of 1,505 upregulated and 2,587 downregulated genes when compared to normal tissue. The crucial next step was to find the overlap between these gene sets. As illustrated in the Venn diagrams this comparison pinpointed a core set of 192 consistently upregulated and 204 consistently downregulated genes ( Figure 3), forming our final list of 396 shared DEGs (sDEGs) for downstream analysis.

**Figure 2.**
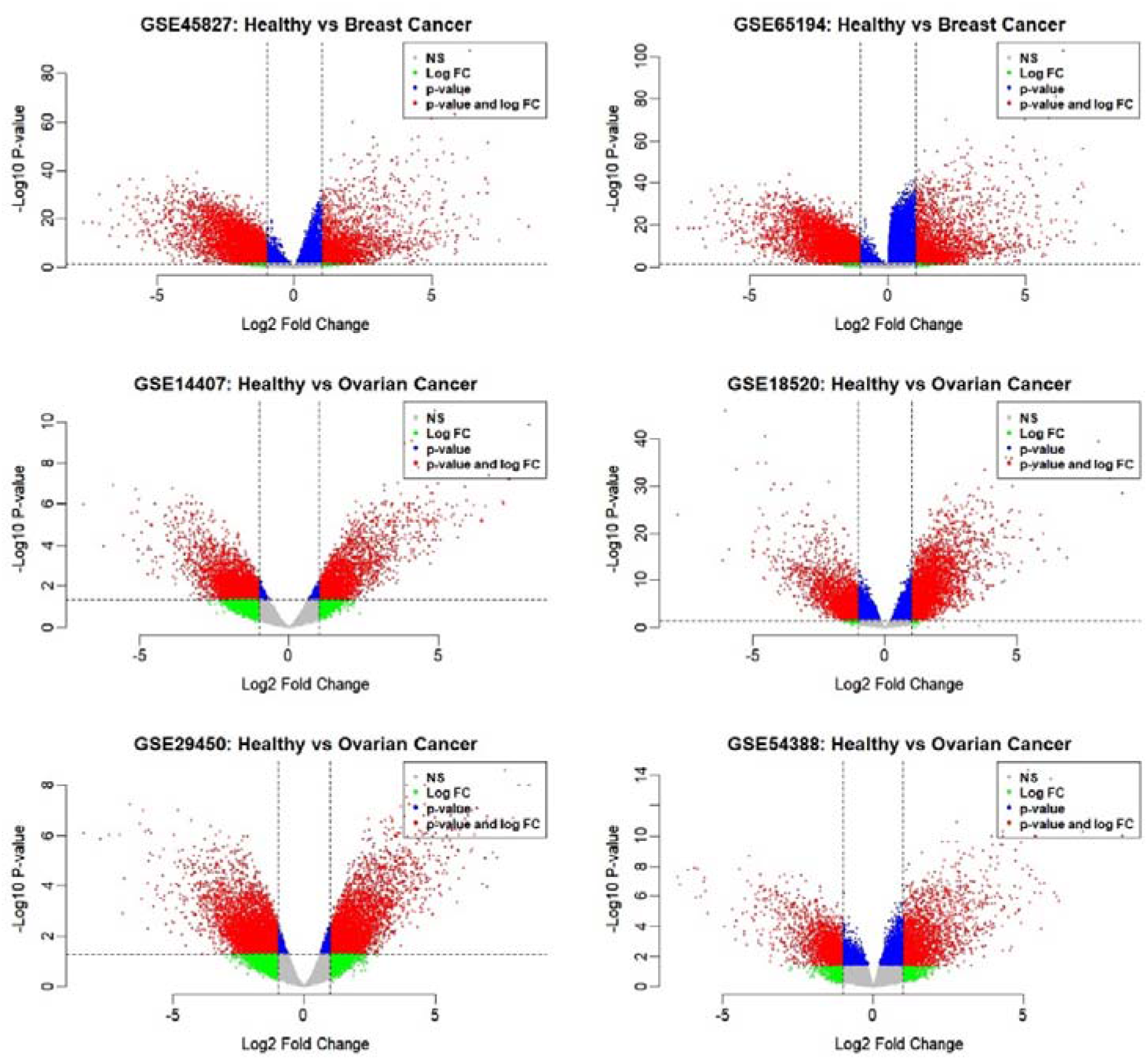
Identification of Differentially Expressed Genes (DEGs) in breast and ovarian cancer. Volcano plots displaying the gene expression profiles for healthy versus breast cancer tissues (top row: GSE45827, GSE65194) and healthy versus ovarian cancer tissues (middle and bottom rows: GSE14407, GSE18520, GSE29450, GSE54388). The x-axis represents the log2 fold change (Log2FC), and the y-axis represents the -log10(p-value). Vertical and horizontal dashed lines indicate the significance thresholds (|log FC| > 1 and adjusted p-value < 0.05, respectively). Red points signify genes that meet both criteria for differential expression.

**Figure 3.**
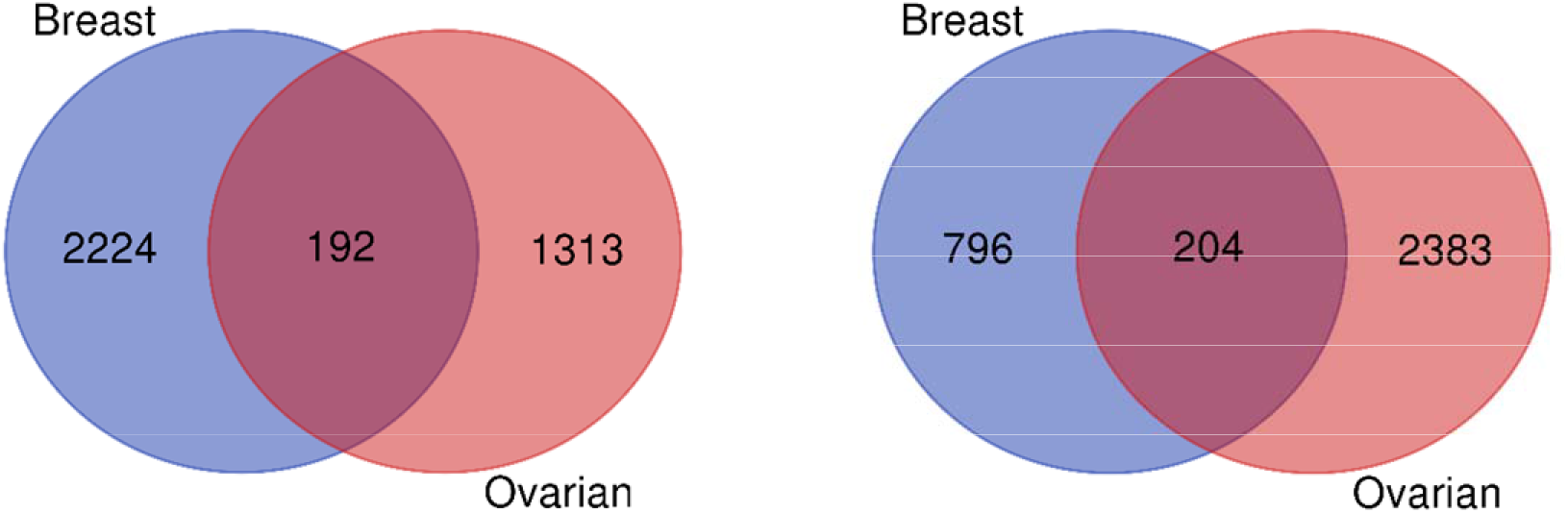
Overlap of significant DEGs between breast and ovarian cancer. Venn diagrams illustrating the shared molecular signature between the two diseases. The left diagram displays the intersection of upregulated genes, identifying 192 shared genes. The right diagram displays the intersection of downregulated genes, revealing 204 shared genes

### 3.2. Identification of shared key genes through PPI network analysis

By applying stringent topological algorithms to our shared DEG PPI network, we distilled a complex transcriptomic profile down to 10 highly connected hub genes (sKGs). The composition of this 10-gene signature (Figure 4) strongly implies that the shared oncogenic drive between breast and ovarian tissues is rooted in severe cell cycle dysregulation combined with the loss of key tumor suppressors.

**Figure 4.**
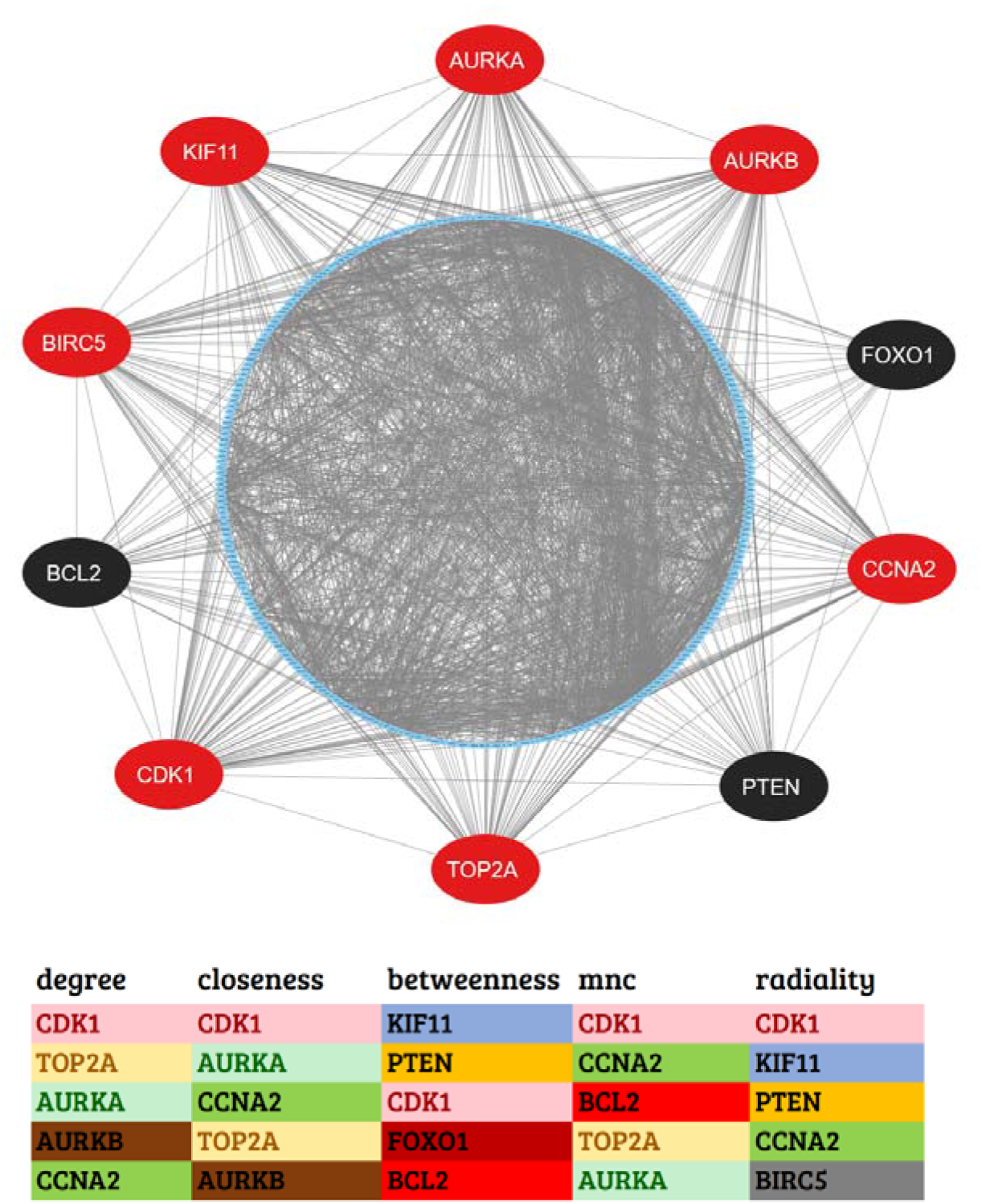
Identification of shared hub genes. (Top) PPI network of the 396 shared DEGs, highlighting the 10 most critical hub genes in the outer ring (red = upregulated; black = downregulated). **(Bottom)** cytoHubba analysis displaying the top five hub genes across five topological algorithms. Colors highlight the overlap of genes across different ranking methods.

CDK1 dominated the topological rankings, acting as the primary bottleneck and most connected node in the shared network. CDK1 is a master regulator of the G2/M phase transition.^101^ Its prominent position, alongside other highly ranked, upregulated mitotic regulators such as CCNA2 (Cyclin A2), TOP2A, and the Aurora kinases (AURKA and AURKB), paints a clear picture of hyperactive cellular proliferation. Both AURKA and AURKB are essential for centrosome maturation and chromosome segregation ^102^; their shared upregulation suggests a common vulnerability to mitotic stress in both malignancies. Therefore, targeting the G2/M checkpoint or Aurora kinase activity may represent a viable dual-therapy approach for both breast and ovarian cohorts.^103^

Conversely, our network analysis identified three critical downregulated nodes: PTEN, FOXO1, and BCL2. PTEN is a classic tumor suppressor and the primary antagonist of the PI3K/AKT signaling pathway,^104^ which is notoriously altered in women’s cancers.^105,106^ The concurrent downregulation of FOXO1, a downstream target often repressed by hyperactive AKT signaling, suggests that the shared oncogenic network heavily relies on the evasion of apoptosis and sustained metabolic survival.^107^

### 3.3. Gene Ontology and KEGG pathway enrichment analysis

The GO Biological Process analysis showed that these hub genes are most significantly associated with “Cell Cycle G2/M Phase Transition,” “Microtubule Cytoskeleton Organization Involved in Mitosis,” and “Mitotic Spindle Organization,” alongside “Regulation of Apoptotic Process” and “DNA Damage Response,” reflecting the mitotic and apoptotic duality of the gene set. In terms of molecular function, the hub genes were enriched for “Protein Heterodimerization Activity,” “Kinase Binding,” and “Protein Serine/Threonine Kinase Activity.” The GO Cellular Component analysis linked the hub genes predominantly to the “Spindle Microtubule,” “Microtubule,” and “Nucleus,” the last of which captured all ten hub genes simultaneously. The KEGG pathway analysis showed that the hub genes converge on “Cellular Senescence,” “Pathways in Cancer,” and the “p53 Signaling Pathway,” directly reflecting the TP53-centred convergence between breast and ovarian carcinoma described earlier (Supplementary Table 1 and Fig. 5). Of 393 Biological Process terms, 52 Molecular Function terms, 32 Cellular Component terms, and 75 KEGG pathways returned, 294, 29, 16, and 29 respectively remained significant after multiple-testing correction (adjusted P < 0.05), indicating that the hub-gene set is not enriched for an isolated function but for a broad, internally consistent proliferative and survival program.

**Figure 5.**
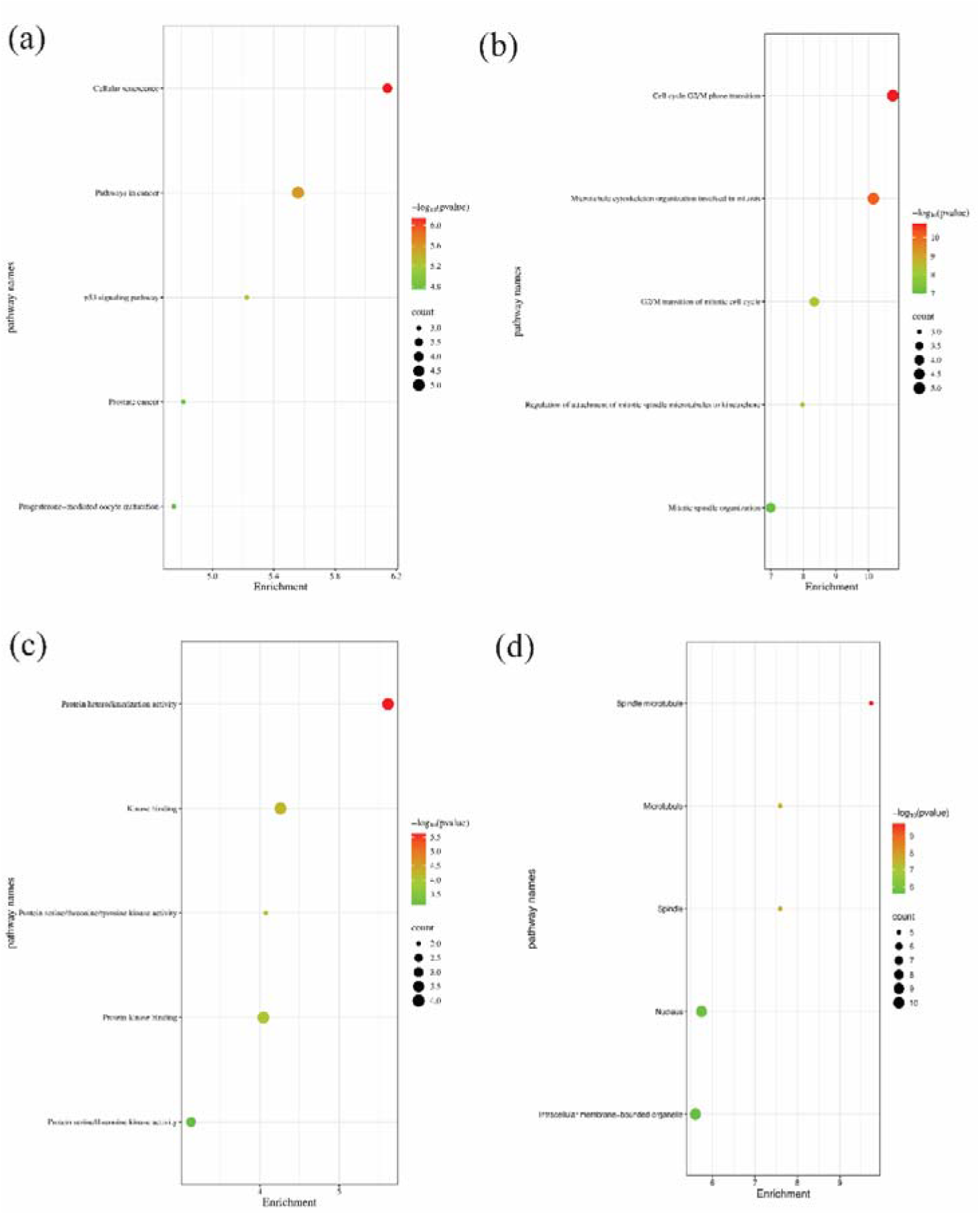
Functional enrichment analysis of hub genes. Bubble charts illustrating the top significantly enriched terms for **(a)** KEGG pathways, **(b)** GO Biological Processes, **(c)** GO Molecular Functions, and **(d)** GO Cellular Components. Bubble size indicates the number of involved genes, and the color gradient represents statistical significance (−log10 *P*-value).

### 3.4. Regulatory network analysis of sKGs

To identify the top-ranking transcription factors (TFs) that act as pivotal transcriptional regulators of the hub genes, we constructed an interaction network between the hub genes and transcription factors. From this network, we determined the five most significant TFs: TP53, STAT3, TFAP2A, FOXC1, and GATA2, based on a degree threshold of ≥ 7. In the visual representation (Fig. 6), these key TFs are depicted as dark-blue hexagons, while hub genes are represented by red and black ellipses.

**Figure 6.**
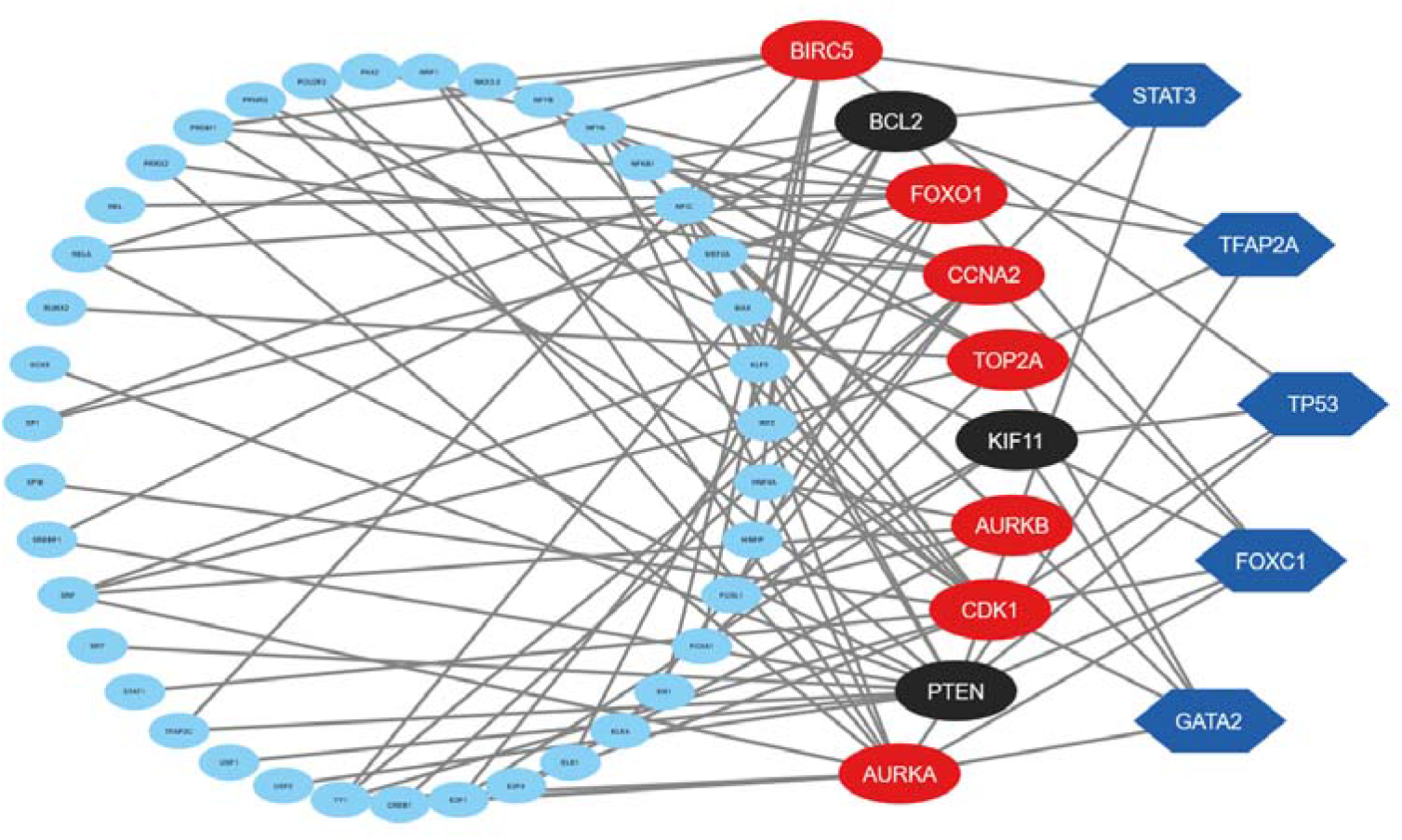
Transcription factor-hub gene interaction network. The interaction network between hub genes and transcription factors (TFs) is shown. Key TFs (TP53, STAT3, TFAP2A, FOXC1, and GATA2) are depicted as dark-blue hexagons, while hub genes are represented by red and black ellipses.

To further identify the primary microRNAs (miRNAs) involved in post-transcriptional regulation of the hub genes, we developed a hub gene–miRNA interaction network. The top five miRNAs hsa-miR-21-5p, hsa-miR-26a-5p, hsa-miR-16-5p, hsa-miR-15a-5p, and hsa-let-7a-5p were identified as critical regulators with a degree of ≥ 8. In the corresponding figure (Fig. 7), these key miRNAs are shown as magenta rectangles, while hub genes are illustrated as red and black ellipses.

**Figure 7.**
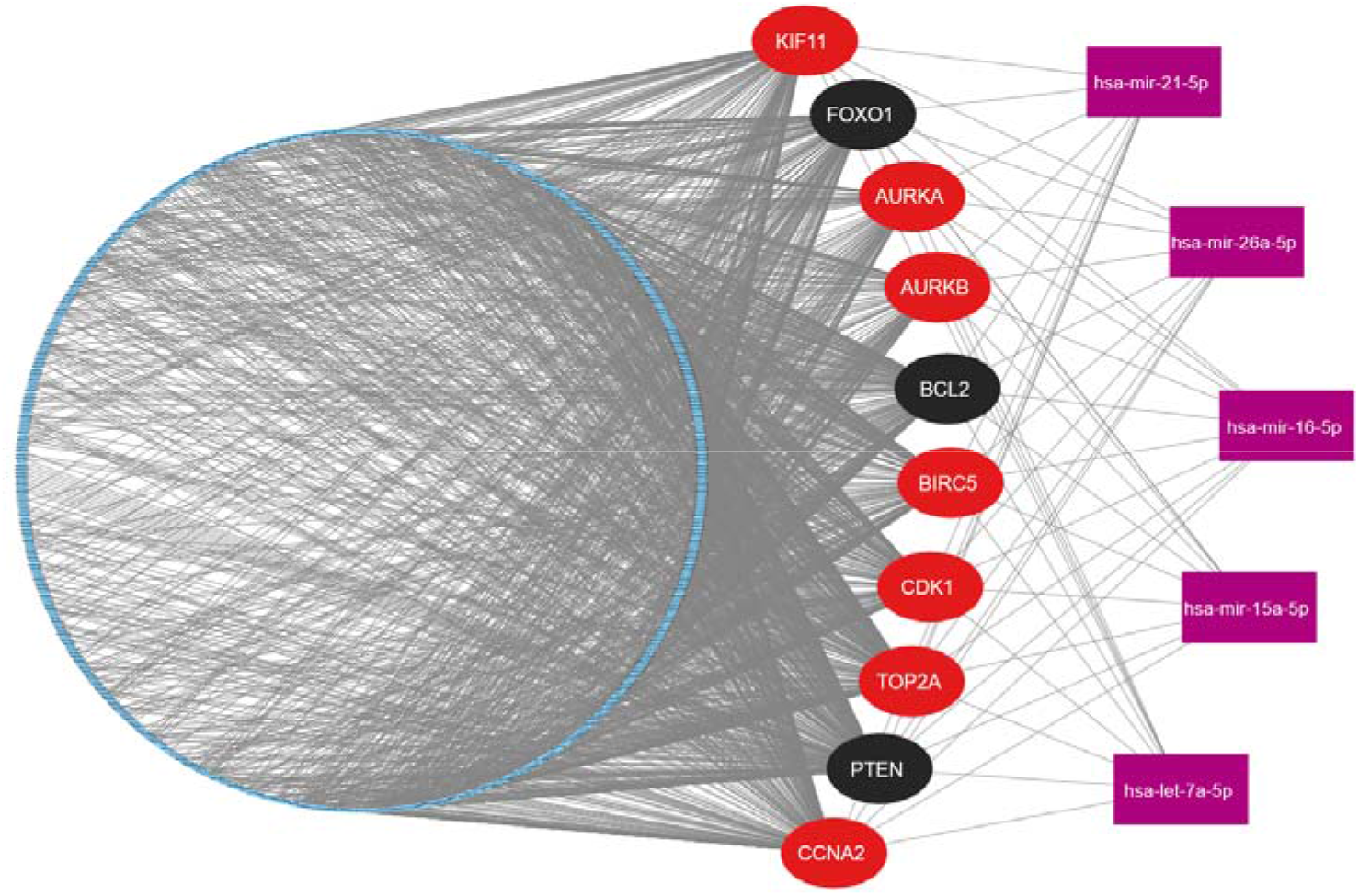
miRNA–hub gene interaction network. The interaction network between hub genes and miRNAs is shown. Key miRNAs (hsa-miR-21-5p, hsa-miR-26a-5p, hsa-miR-16-5p, hsa-miR-15a-5p and hsa-let-7a-5p) are shown as magenta rectangles, while hub genes are represented by red and black ellipses.

### 3.5. Gene expression analysis

In the GEPIA2 database, transcriptional expression differences of the ten hub genes between tumor and normal tissue were validated in both breast (BRCA, tumor n = 1085, normal n = 291) and ovarian (OV, tumor n = 426, normal n = 88) cohorts (Fig. 8). Of the ten hub genes, seven — AURKA, AURKB, BIRC5, CCNA2, CDK1, KIF11, and TOP2A were significantly up-regulated (p < 0.05) in tumor relative to normal tissue in both BRCA and OV (Fig. 8). FOXO1 was significantly down-regulated in both BRCA and OV, in line with its role as a tumor-suppressive node of the PI3K–AKT–FOXO axis. PTEN showed no significant difference between tumor and normal tissue in either BRCA or OV, despite its established role as a negative regulator of PI3K–AKT signalling. BCL2 showed a directional inconsistency between the two cancers, underscoring that its role in apoptotic regulation is context-dependent rather than uniform across breast and ovarian tissue. This inconsistency is reported rather than concealed, consistent with the directional heterogeneity anticipated for BIRC5-family and FOXO-axis genes at the outset of this study.

**Figure 8.**
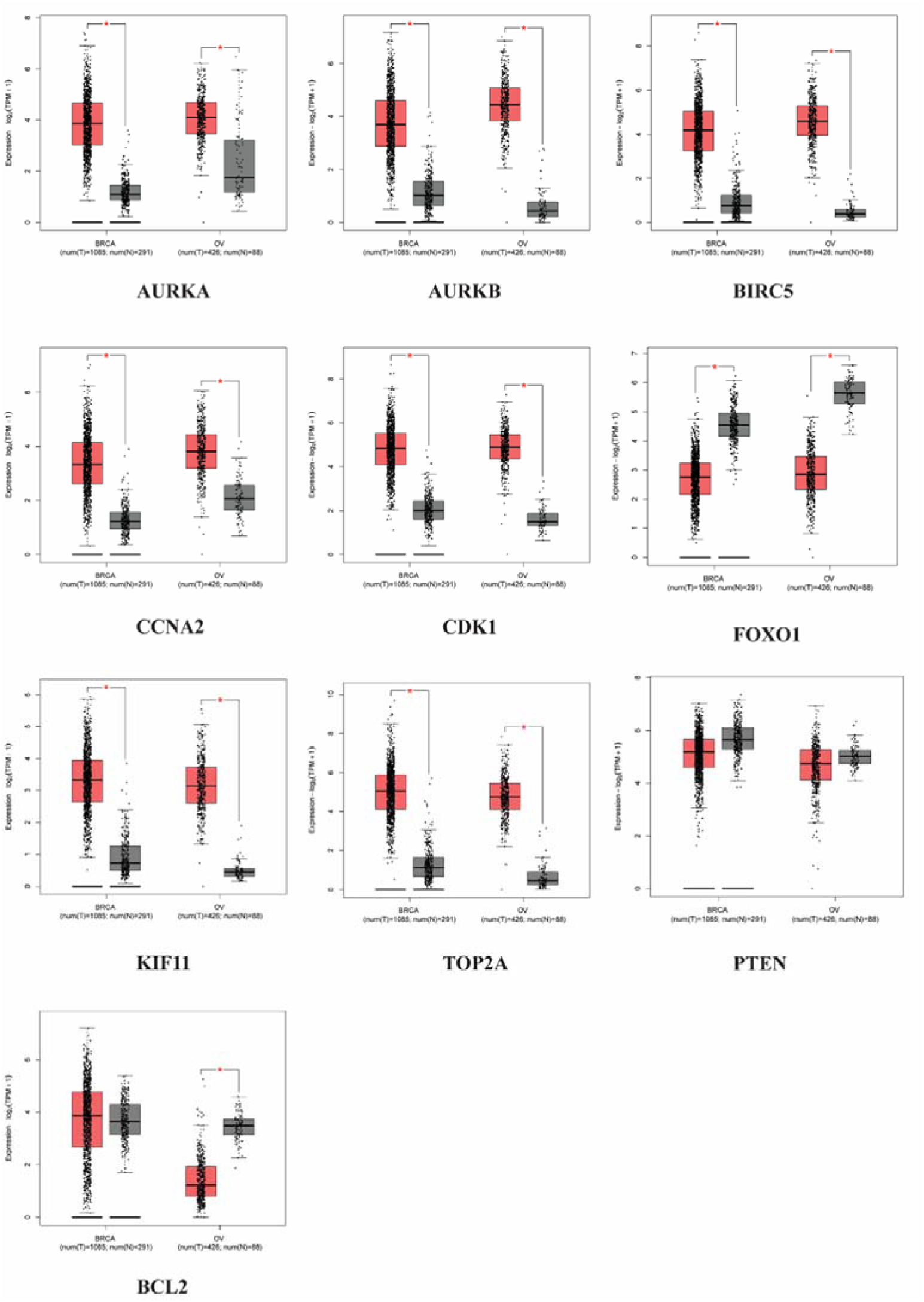
GEPIA2 validation of hub-gene expression in breast (BRCA) and ovarian (OV) cancers. Boxplots show tumor–normal expression differences for ten hub genes. Seven genes were significantly upregulated in both cancers, FOXO1 was downregulated, PTEN showed no significant difference, and BCL2 exhibited a cancer-specific expression pattern.

### 3.6. Virtual screening of phytochemicals

#### 3.6.1. Physicochemical-based screening

An initial library of 70,478 phytochemical compounds was retrieved from FooDB. Structures were standardized in RDKit and filtered against the Lipinski, Ghose, Veber, Egan, and Muegge drug-likeness rule sets, with compounds flagged by PAINS, Brenk, or NIH structural alerts subsequently excluded, reducing the library to 4,190 compounds (5.95% of the initial set).

#### 3.6.2. Machine learning-based screening

The individual target models demonstrated moderate-to-good predictive performance, with R² values ranging from 0.50 to 0.77 and RMSE values from 0.35 to 0.40. AURKA showed the strongest performance (R² = 0.77, RMSE = 0.35), followed by CDK1 (R² = 0.74, RMSE = 0.36) and TOP2A (R² = 0.71, RMSE = 0.37). In contrast, FOXO1 (R² = 0.50, RMSE = 0.40) and CCNA2 (R² = 0.54, RMSE = 0.40) showed comparatively weaker performance. The AURKB, KIF11, and BIRC5 models achieved R² values of 0.69, 0.61, and 0.58, respectively. Notably, combining the bioactivity datasets from all eight proteins improved overall performance, with the pooled multi-target model achieving an R² of 0.79 and an RMSE of 0.28. This improvement may reflect the larger and more chemically diverse training dataset, enabling the model to learn broader molecular patterns associated with biological activity. The compounds were then screened using the bioactivity prediction (ML/QSAR) model and 862 compounds with a predicted pIC50 > 5 were retained (20.60% of the drug-likeness-filtered set) (Supplementary file 1).

#### 3.6.3. Pharmacokinetic and toxicity screening

The remaining 862 compounds were evaluated in Deep-PK for AMES mutagenicity, carcinogenicity, and drug-induced liver injury (DILI); compounds predicted positive for any of the three endpoints were excluded, yielding a final ligand library of 233 compounds carried forward to molecular docking (Supplementary file 2).

### 3.7. Molecular docking analysis

Molecular docking was performed against eight of the ten hub gene targets — AURKA, AURKB, BIRC5, CCNA2, CDK1, FOXO1, KIF11, and TOP2A. PTEN and BCL2 were excluded from the docking arm: PTEN showed no significant differential expression in either BRCA or OV in the GEPIA2 validation and was retained only as a regulatory/prognostic biomarker, while BCL2 showed a discordant expression direction between BRCA and OV in the same validation, precluding a consistent oncogenic-hub rationale for structure-based screening. FOXO1 was retained in the docking arm, but given its role as a consistently down-regulated tumor suppressor in both cancers, compounds against this target were evaluated as candidate modulators/reactivators of FOXO1 activity rather than inhibitors.

Compounds were ranked by predicted binding free energy (ΔG, kcal/mol), and a distinct lead compound was assigned to each target, avoiding reuse of a compound already claimed by a higher-affinity target elsewhere in the panel, (Fig. 9).

**Figure 9.**
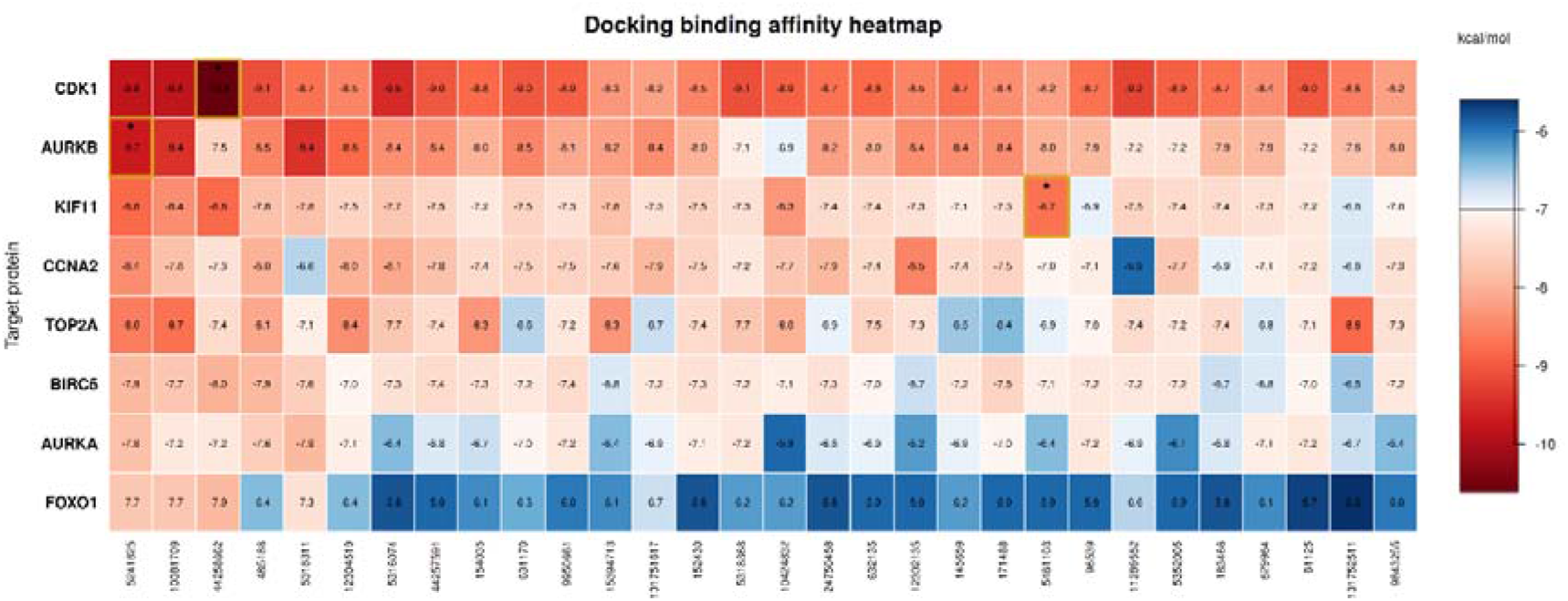
Docking binding-affinity heatmap of the top 30 ranked compounds against eight hub-gene targets. Red = strong binding (≤ −7.0 kcal/mol); blue = weaker binding. Gold-outlined, starred cells mark three representative high-affinity ligand–target pairs

Across the three complexes for which full interaction profiles were resolved, binding was governed predominantly by hydrophobic contacts, with each ligand additionally anchored by one or two directional interactions. At AURKB, CID 5241825 (−9.7 kcal/mol) is held in place by a hydrogen bond with Phe219, with the surrounding pocket dominated by alkyl and pi-alkyl contacts centered on Val91, Leu207, and Phe88. At CDK1, CID 44258662 (−10.7 kcal/mol) forms the densest and most energetically favorable network of the three, with overlapping pi-sigma and alkyl contacts spanning Phe80, Ile10, Val18, and Leu135, in line with deep occupation of the ATP-binding cleft. At KIF11, CID 5461103 (−8.5 kcal/mol) breaks from this hydrophobic-dominant pattern: the complex is instead stabilized by a carbon-hydrogen bond and a pi-anion interaction with Glu116, pointing to a more polar binding environment than the other targets. Full residue-level contacts, bond types, and interaction distances for all three complexes are reported in Table 2 (Fig. 10)

**Figure 10.**
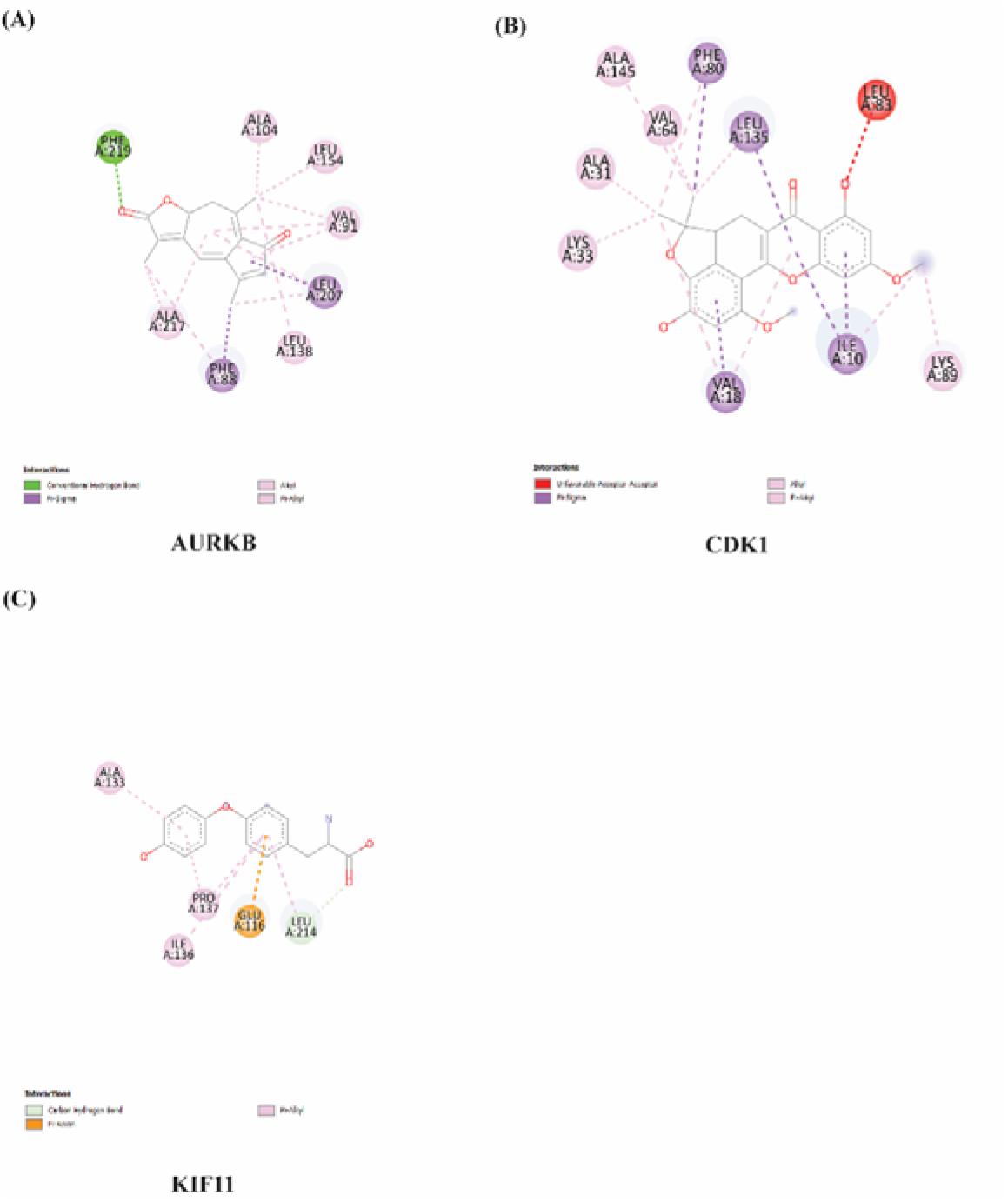
2D binding interaction profiles of prioritized docking complexes. Visualizations showing non-covalent interactions (such as hydrogen bonds, alkyl/pi-alkyl contacts, and pi-sigma interactions) between small molecule ligands and key residue active sites for: (A) AURKB + CID 5241825, (B) CDK1 + CID 44258662 and (C) KIF11 + CID 5461103.

**Table 1.** Summary of Gene Expression Omnibus (GEO) datasets utilized in the analysis, including accession numbers, country of origin, publication/release year, microarray platform, organism, and the number of case and control samples.

| **GEO datasets** | **Country** | **Year** | **Platform** | **Organism** | **Cases** | **Control** |
|---|---|---|---|---|---|---|
| GSE45827 | France | 2016 | GPL570 [HG-U133_Plus_2] Affymetrix Human Genome U133 Plus 2.0 Array | *Homo sapiens* | 130 | 25 |
| GSE65194 | France | 2015 | GPL570 [HG-U133_Plus_2] Affymetrix Human Genome U133 Plus 2.0 Array | *Homo sapiens* | 167 | 11 |
| GSE14407 | USA | 2009 | GPL570 [HG-U133_Plus_2] Affymetrix Human Genome U133 Plus 2.0 Array | *Homo sapiens* | 12 | 12 |
| GSE18520 | USA | 2009 | GPL570 [HG-U133_Plus_2] Affymetrix Human Genome U133 Plus 2.0 Array | *Homo sapiens* | 53 | 10 |
| GSE29450 | USA | 2011 | GPL570 [HG-U133_Plus_2] Affymetrix Human Genome U133 Plus 2.0 Array | Homo sapiens | 10 | 5 |
| GSE54388 | USA | 2014 | GPL570 [HG-U133_Plus_2] Affymetrix Human Genome U133 Plus 2.0 Array | Homo sapiens | 8 | 6 |

**Table 2.** Binding interactions of prioritized docking complexes

| Complex | Binding energy (kcal/mol) | Residue | Bond type | Distance (Å) |
|---|---|---|---|---|
| AURKB + CID 5241825 | −9.7 | Phe219 | H-bond | 1.98 |
|  |  | Leu207 | Pi-Sigma | 3.88 |
|  |  | Phe88 | Pi-Sigma | 3.54 |
|  |  | Val91 | Alkyl | 5.4 |
|  |  | Ala104 | Alkyl | 3.69 |
|  |  | Leu207 | Alkyl | 5.2 |
|  |  | Ala217 | Alkyl | 4.24 |
|  |  | Ala217 | Alkyl | 4.29 |
|  |  | Val91 | Alkyl | 5.23 |
|  |  | Leu138 | Alkyl | 4.88 |
|  |  | Leu154 | Alkyl | 4.19 |
|  |  | Leu207 | Alkyl | 4.64 |
|  |  | Phe88 | Pi-Alkyl | 4.86 |
|  |  | Val91 | Pi-Alkyl | 4.93 |
| CDK1 + CID 44258662 | −10.7 | Phe80 | Pi-Sigma | 3.79 |
|  |  | Ile10 | Pi-Sigma | 3.96 |
|  |  | Ile10 | Pi-Sigma | 3.77 |
|  |  | Val18 | Pi-Sigma | 3.56 |
|  |  | Leu135 | Pi-Sigma | 3.76 |
|  |  | Val18 | Alkyl | 3.96 |
|  |  | Ala31 | Alkyl | 4.4 |
|  |  | Lys33 | Alkyl | 4.41 |
|  |  | Val64 | Alkyl | 3.93 |
|  |  | Leu135 | Alkyl | 5.42 |
|  |  | Ala145 | Alkyl | 3.73 |
|  |  | Ile10 | Alkyl | 5.08 |
|  |  | Lys89 | Alkyl | 4.66 |
|  |  | Val18 | Pi-Alkyl | 5.08 |
|  |  | Phe80 | Pi-Alkyl | 4.19 |
| KIF11 + CID 5461103 | −8.5 | Leu214 | C-H bond | 3.66 |
|  |  | Glu116 | Pi-Anion | 3.94 |
|  |  | Ile136 | Pi-Alkyl | 5.34 |
|  |  | Pro137 | Pi-Alkyl | 5.21 |
|  |  | Leu214 | Pi-Alkyl | 4.14 |
|  |  | Ala133 | Pi-Alkyl | 4.67 |
|  |  | Pro137 | Pi-Alkyl | 4.27 |

### 3.8. Docking validation

Decoy-based validation showed target-dependent discrimination between the reference ligands and their corresponding property-matched decoys. The AURKB reference ligand (−9.7 kcal/mol) showed the strongest separation, with a Z-score of −1.77 (Table 3) and no decoy achieving a more favorable docking score, placing the ligand at the 98th percentile of the decoy distribution. Similarly, the CDK1 reference ligand (−10.7 kcal/mol) showed favorable ranking relative to the decoys (Z-score = −1.50), with no decoy outperforming it and a 94th-percentile ranking. In contrast, discrimination was weaker for KIF11. Ten of 50 KIF11 decoys (20%) achieved a more favorable docking score than the reference ligand, corresponding to a 78th-percentile ranking and a Z-score of −0.77. Overall, the decoy analysis provided stronger support for the docking-based prioritization of the AURKB and CDK1 ligands, while the results for KIF11 indicated comparatively greater overlap between reference-ligand and decoy docking scores.

**Table 3.** Decoy-based validation of molecular docking performance for the three target proteins.

| **Target** | **Reference Ligand Score (kcal/mol)** | **Decoy Mean ± SD (kcal/mol)** | **Z-score** | **Best Decoy Score (kcal/mol)** | **Decoys Outperforming Reference** | **Decoy Percentile** |
|---|---|---|---|---|---|---|
| **AURKB** | −9.7 | −7.93 ± 1.00 | −1.77 | −9.7 | 0/50 (0%) | 98th |
| **CDK1** | −10.7 | −9.19 ± 1.01 | −1.50 | −10.7 | 0/50 (0%) | 94th |
| **KIF11** | −8.5 | −7.73 ± 1.00 | −0.77 | −9.4 | 10/50 (20%) | 78th |

#### 3.8.1. Redocking of co-crystallised ligands

All three co-crystallised ligands returned to their experimental binding mode (Table 4; Figure 11). The top-ranked pose of LZ9 in CDK1 deviated from the crystallographic pose by 0.66 Å, that of VX6 in AURKB by 0.94 Å and that of K5E in KIF11 by 0.42 Å, all below the 2.0 Å criterion, and in each case the top-ranked pose was also the closest of the nine poses to the crystal structure. The scoring function therefore identifies the experimental pose as the most favourable one at all three targets when the location of the site is supplied.

**Figure 11.**
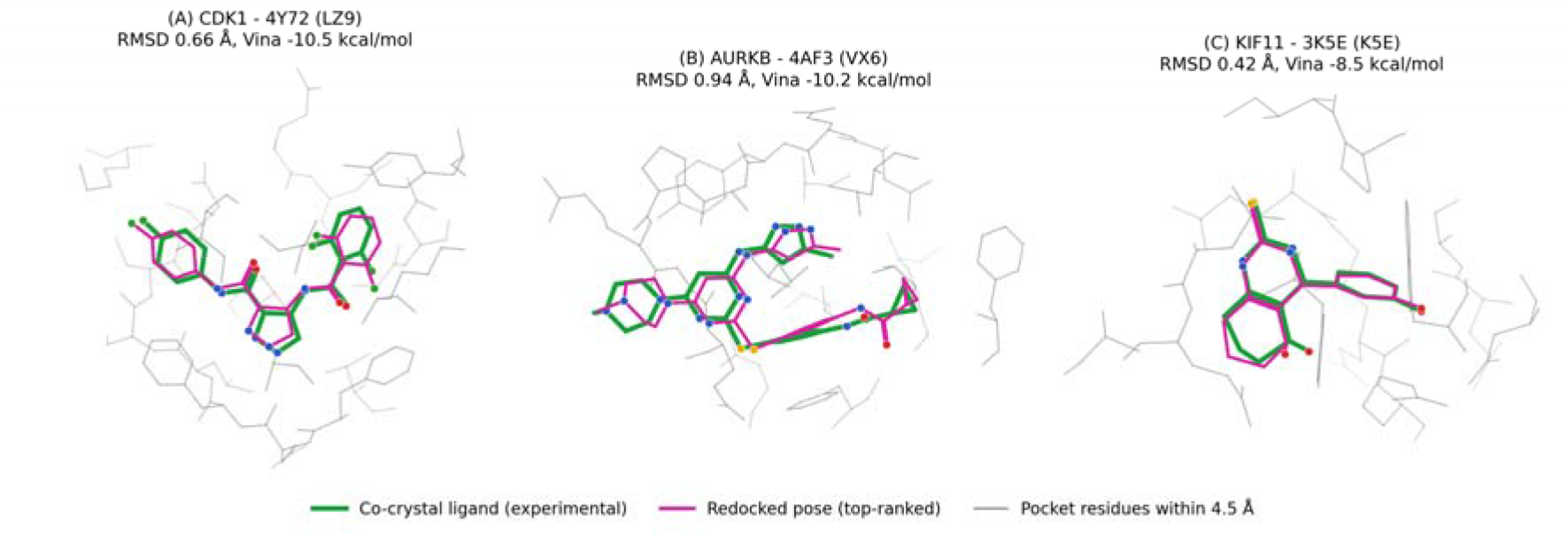
Redocking of co-crystallised ligands in the site-centred box (22 Å, exhaustiveness 32). Experimental pose (green) and top-ranked redocked pose (magenta) of (A) LZ9 in CDK1 (PDB 4Y72), (B) VX6 in AURKB (PDB 4AF3) and (C) K5E in KIF11 (PDB 3K5E). Residues within 4.5 Å of the crystallographic ligand are shown as grey lines. Heavy-atom RMSD and Vina score of the top-ranked pose are given above each panel.

**Table 4.** Redocking of co-crystallised ligands. Site-centred box: 22 Å on the crystallographic ligand, exhaustiveness 32. Whole-protein box: receptor bounding box plus 5 Å, exhaustiveness 16/32/64 × 3 seeds (nine runs). Site recovery, centroid of the top-ranked pose within 5.0 Å of the crystallographic ligand; pose accuracy, RMSD < 2.0 Å.

| **Target** | **PDB** | **Co-crystal ligand** | **RMSD (Å)** | **Whole-protein: site recovered** |
|---|---|---|---|---|
| **CDK1** | 4Y72 | LZ9 | 0.66 | 9/9 |
| **AURKB** | 4AF3 | VX6 | 0.94 | 3/9 |
| **KIF11** | 3K5E | K5E | 0.42 | 9/9 |

#### 3.8.2. Benchmarking against known inhibitors

Docking scores separated known actives from measured inactives only modestly (Table 5). The ROC-AUC was 0.566 (95% CI 0.44–0.69) for CDK1, 0.687 (0.57–0.80) for AURKB and 0.724 (0.60–0.83) for KIF11; the interval for CDK1 includes 0.5. The AUC obtained from heavy-atom count alone was 0.617, 0.731 and 0.676, respectively, equal to or higher than that of the docking score for CDK1 and AURKB and slightly lower for KIF11. In this benchmark the scores therefore mainly track ligand size, and they are better read as a coarse filter than as a predictor of affinity. Median scores of actives and inactives were −9.82 and −9.68 kcal/mol for CDK1, −9.72 and −9.12 for AURKB, and −9.23 and −8.43 for KIF11. The inactives are compounds with measured weak activity, not property-matched decoys that are merely presumed inactive, and the sets are small (40 per class), so the confidence intervals are wide.

**Table 5.** Docking benchmark against measured ChEMBL actives (pChEMBL ≥ 7) and inactives (pChEMBL ≤ 5). AUC, area under the ROC curve (95% bootstrap CI); size-only AUC, AUC obtained from heavy-atom count alone.

| **Target** | **ChEMBL target** | **Actives / inactives** | **Median Vina, actives** | **Median Vina, inactives** | **ROC-AUC (95% CI)** | **Size-only AUC** |
|---|---|---|---|---|---|---|
| **CDK1** | CHEMBL308 | 40 / 40 | −9.82 | −9.68 | 0.566 (0.44–0.69) | 0.617 |
| **AURKB** | CHEMBL2185 | 40 / 40 | −9.72 | −9.12 | 0.687 (0.57–0.80) | 0.731 |
| **KIF11** | CHEMBL4581 | 40 / 40 | −9.23 | −8.43 | 0.724 (0.60–0.83) | 0.676 |

Placed on this scale (Table 6), artonin L scored −10.7 kcal/mol at CDK1, better than 80% of the known CDK1 actives and 87.5% of the measured inactives, and was the only one of the three ligands to fall in the upper range of the measured actives of its own target. Taraxacin scored −9.1 kcal/mol at AURKB, out-scoring 30% of the known AURKB actives and 47.5% of the inactives; normalised per heavy atom it out-scored 100% of the actives, reflecting its small size (18 heavy atoms against a median of 33 for the AURKB actives). L-Thyronine at KIF11 (−8.9 kcal/mol) out-scored 30% of the actives and 72.5% of the inactives, the same share of actives as taraxacin at AURKB, and likewise reached 100% per heavy atom. Scores were also not target-selective: the three ligands scored between −7.1 and −10.6 kcal/mol at the two non-cognate targets, and L-thyronine scored −8.86 kcal/mol at CDK1 and −8.85 kcal/mol at its own target KIF11. The benchmark therefore places artonin L at CDK1 in the upper range of measured CDK1 inhibitors, while taraxacin and L-thyronine fall in the lower part of the range at their own targets on raw score and at the top of it once normalised for ligand size.

**Table 6.** Position of the three prioritised ligands relative to the benchmark actives and inactives of each target. “Out-scored” is the percentage of benchmark compounds of that target with a less favourable Vina score. Cognate pairs are listed first.

| **Target** | **Ligand** | **Heavy atoms** | **Vina (kcal/mol)** | **Actives out-scored (%)** | **Inactives out-scored (%)** |
|---|---|---|---|---|---|
| **CDK1** | Artonin L (own target) | 29 | −10.7 | 80 | 87.5 |
| **CDK1** | Taraxacin | 18 | −10.6 | 75 | 82.5 |
| **CDK1** | L-Thyronine | 20 | −8.9 | 20 | 25.0 |
| **AURKB** | Taraxacin (own target) | 18 | −9.1 | 30 | 47.5 |
| **AURKB** | Artonin L | 29 | −8.7 | 18 | 27.5 |
| **AURKB** | L-Thyronine | 20 | −8.3 | 2 | 22.5 |
| **KIF11** | L-Thyronine (own target) | 20 | −8.9 | 30 | 72.5 |
| **KIF11** | Artonin L | 29 | −7.1 | 0 | 7.5 |
| **KIF11** | Taraxacin | 18 | −7.5 | 2 | 17.5 |

### 3.9. Molecular dynamics simulation

#### 3.9.1. Root-mean-square deviation (RMSD)

RMSD trajectories were analyzed to evaluate how effectively the candidate ligand maintains structural stability across the three target complexes. Upon binding, the KIF11 complex rapidly reached equilibrium within the first 10 ns, maintaining a steady plateau between 2.5 and 3.2 Å. CDK1 exhibited a controlled, two-phase dynamic profile, remaining stable at 2.0–2.5 Å for the initial 40 ns before undergoing a slight structural shift to establish a second plateau at 2.7–3.5 Å for the remainder of the simulation (Fig 12a).

**Figure 12.**
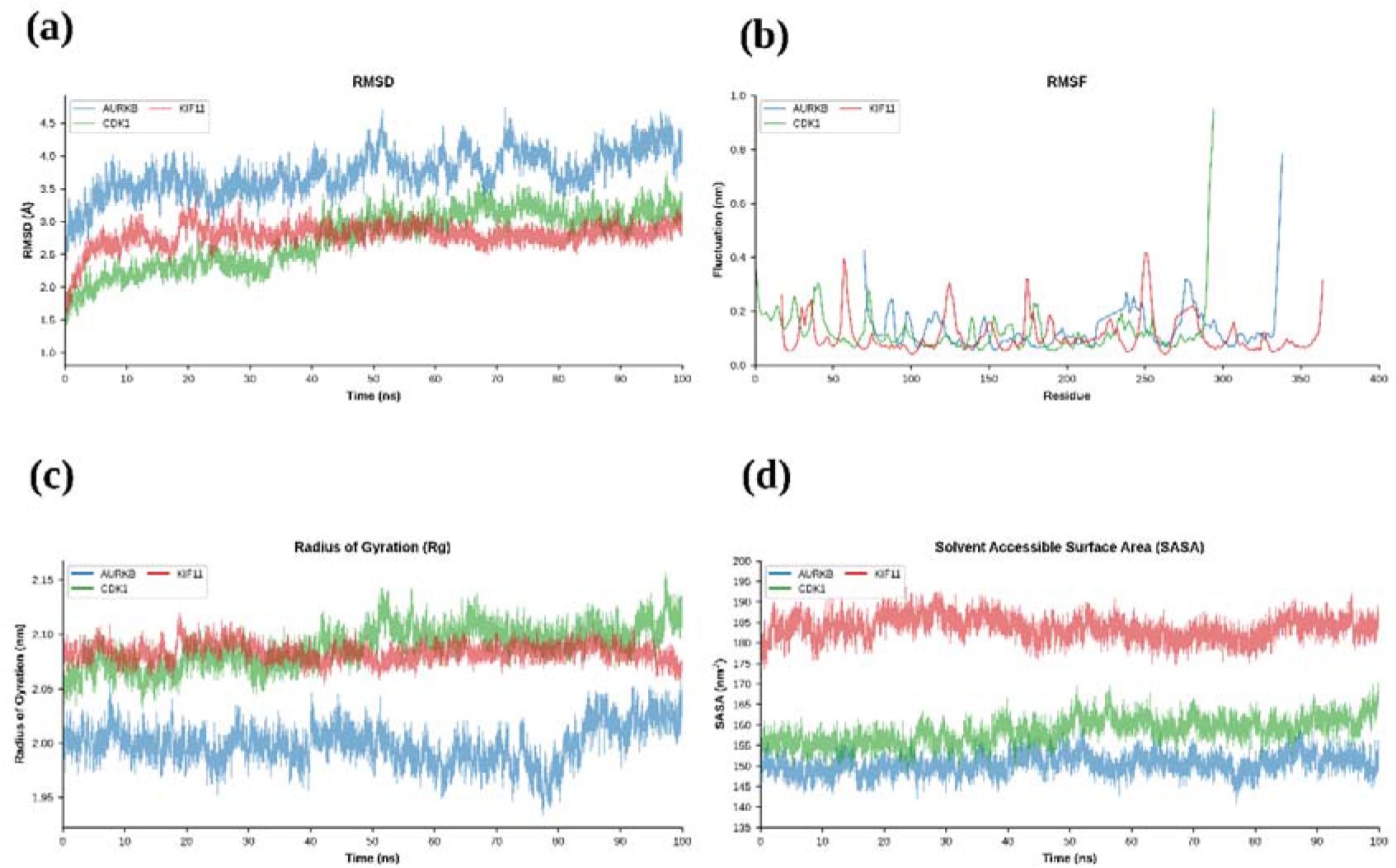
Molecular dynamics simulation analysis of the three protein–ligand complexes over a 100 ns trajectory. (a) Root-mean-square deviation (RMSD) of the protein backbones as a function of simulation time, (b) Residue-wise root-mean-square fluctuation (RMSF), (c) radius of gyration (Rg), and (d) solvent-accessible surface area (SASA).

**Figure 13.**
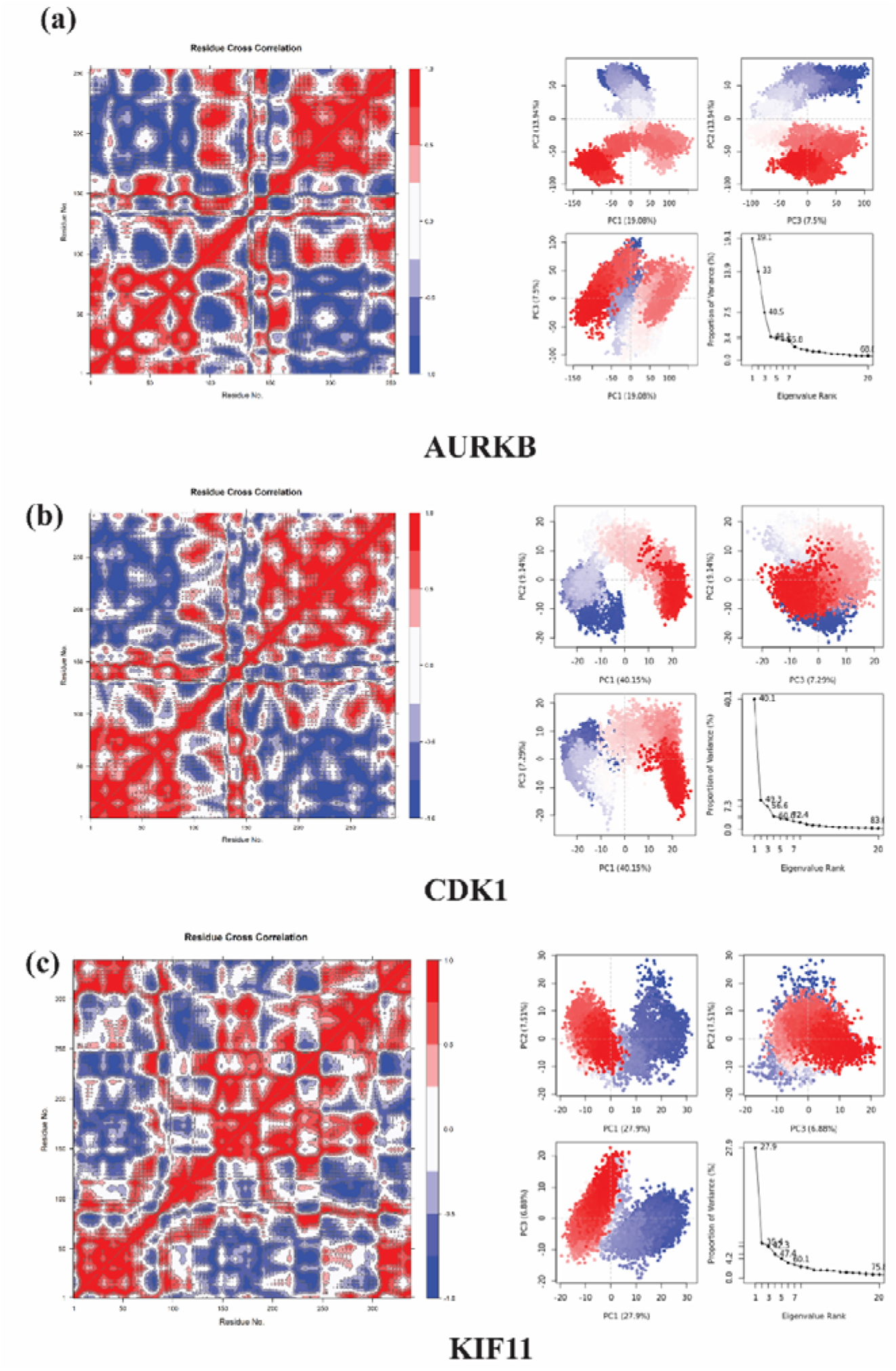
Residue cross-correlation matrices and principal component analysis (PCA) of the molecular dynamics trajectories of the protein–ligand complexes. (a–c) AURKB, CDK1, and KIF11, respectively. Left panels show residue cross-correlation matrices, with red and blue regions indicating positively correlated and anticorrelated residue motions, respectively, and white regions representing weak or negligible correlation. Right panels show PCA projections along the first three principal components (PC1, PC2, and PC3) and the corresponding eigenvalue spectra illustrating the distribution of positional fluctuations across the principal components.

In contrast, the AURKB complex displayed greater structural flexibility. AURKB exhibited the largest overall deviations. Its bound structure fluctuated between 3.0 and 4.3 Å for the first 50 ns and subsequently sustained high-amplitude movements ranging from 3.5 to 4.7 Å.

#### 3.9.2. Root-mean-square fluctuation (RMSF)

Root Mean Square Fluctuation (RMSF) is widely monitored to evaluate the dynamic variability and local structural flexibility of individual amino acid residues upon ligand binding.^109^ Based on this principle, RMSF profiles were evaluated throughout the simulations for the three distinct target proteins in this study. Across all complexes, the ligand successfully stabilized the core regions, maintaining baseline fluctuations below 0.1 nm (Fig .12b).

For the KIF11 complex, the ligand maintained a highly rigid core. Localized flexibility was strictly confined to specific surface loops near residues 58, 128, 185, and 252, with maximum fluctuations reaching only 0.4 nm. This suggests that the ligand effectively anchors the protein while accommodating necessary functional loop movements.

The ligand similarly stabilized the central domains of CDK1. Although the N-terminal regions (residues 25–45) exhibited minor flexibility of up to 0.35 nm, the most pronounced movements occurred at their unstructured C-terminal ends near residue 295, peaking at 0.95 nm and 1.05 nm, respectively. These elevated values reflect the inherent mobility of terminal tails rather than any disruption to the ligand-bound core.

A comparable pattern emerged for AURKB. The ligand kept internal loop deviations to a minimum, displaying minor peaks of 0.42 nm near residue 70 and 0.32 nm near residue 285. Additionally, its C-terminal tail exhibited a characteriztic fluctuation spike of 0.78 nm at residue 338.

#### 3.9.3. Solvent-accessible surface area (SASA)

The Solvent Accessible Surface Area (SASA) is widely monitored to evaluate the extent of a candidate ligand’s exposure to solvent molecules within a binding pocket. ^109^ Based on this principle, SASA profiles were evaluated throughout the simulations for the three target proteins in this study. Throughout the simulations, all three complexes exhibited remarkably stable and consistent SASA profiles, devoid of drastic fluctuations. This steady state behavior indicates that the ligand rapidly achieved and maintained a stable binding conformation early in the simulation, without undergoing major spatial reorientations.^110^

Among the targets, the ligand bound AURKB complex displayed the lowest overall SASA values, strictly ranging between 145 and 155 nm² (Fig. 12d). This reduced solvent exposure suggests that the ligand is deeply buried within a highly enclosed binding pocket, effectively shielded from the surrounding aqueous environment. The CDK1 complex maintained slightly higher but equally stable profiles, with SASA value fluctuating narrowly between 150 and 168 nm². This reflects a moderately open yet highly stable interaction.

In contrast, the KIF11 complex exhibited the highest overall solvent exposure, establishing a steady plateau between 175 and 192 nm². Although this points to a more solvent exposed binding mode, potentially indicating that the ligand occupies a broader surface groove or shallower cleft, the lack of sharp deviations confirms that the interaction remained highly stable over time.

Overall, while the burial depth varies depending on the specific architecture of each target protein, ranging from highly shielded in AURKB to partially accessible in KIF11, the consistently steady profiles confirm that the candidate ligand effectively anchors itself and remains structurally robust across all conditions.

#### 3.9.4. Radius of gyration (Rg)

The radius of gyration (Rg) was calculated to monitor the overall compactness and structural dimensions of the three target proteins, as it effectively measures spatial stability and folding behaviors upon ligand binding.^109^ Among the simulated systems, the AURKB complex exhibited the lowest and most stable Rg values, remaining strictly between 1.95 and 2.05 nm (Fig. 12c). This suggests that the ligand induces a highly compact and tightly packed conformation, effectively preventing major unfolding events.^111^

In contrast, KIF11 maintained moderately higher yet stable Rg profile. KIF11 fluctuated narrowly between 2.05 and 2.12 nm. Notably, the CDK1 complex displayed a distinct structural adjustment. Although it initially remained within a moderate range, its Rg profile exhibited a steady upward trend during the latter half of the simulation, gradually increasing to reach a peak of 2.15 nm. This gradual expansion implies that ligand binding leads to a slight increase in the overall dimensions of CDK1, potentially causing certain flexible domains to extend outward.

#### 3.9.5. Principal component analysis (PCA)

Principal Component Analysis (PCA) was performed to characterize the essential dynamics and conformational phase space of the target complexes.^109^ The 2D projections and eigenvalue distributions reveal that the candidate ligand imposes the most rigid restrictions on the CDK1 and KIF11 targets. Specifically, CDK1 displays a highly compact cluster where PC1 accounts for a dominant 40.15% of the variance, capturing 83.0% across the top 20 modes. Similarly, KIF11 exhibits a tightly bounded phase space (PC1: 27.90%). Together, these profiles indicate highly stable and restricted dynamic states that effectively suppress major structural unfolding.^109^

Conversely, AURKB retains greater conformational freedom. The AURKB complex transitions between specific, localized sub-clusters (PC1: 19.08%, top 20: 68.6%), suggesting the stabilization of distinct structural sub-states rather than a singular rigid posture.

#### 3.9.6. Dynamic cross-correlation matrix (DCCM)

Dynamic Cross-Correlation Matrix (DCCM) analysis was performed to evaluate intra-molecular communication and domain-level coordinated motions within the three target proteins upon ligand binding. In the resulting heatmaps, positive correlations (red) represent concerted motions occurring in the same direction, while negative correlations (blue) indicate anti-correlated, opposing structural movements.^112^

The DCCM profiles reveal that the candidate ligand induces strong, highly coordinated global dynamic networks within the AURKB, CDK1, and KIF11 complexes. Both AURKB and CDK1 display large, distinct off-diagonal blocks of intense positive and negative correlations. This signature points to strong structural coupling, where distinct domains move collectively as rigid bodies in a highly concerted or tightly anti-correlated manner. Similarly, KIF11 demonstrates a dense and intricate pattern of intense red and blue regions across its entire sequence. This reflects a tightly coupled dynamic network and robust, long-range allosteric communication throughout the protein architecture.^112^

### 3.10. Binding free-energy calculations (MM/GBSA)

Binding free energies from MM/GBSA differed markedly between the three complexes (Table 8; Figure 14). CDK1–artonin L had the most favourable ΔG_bind_, −31.1 ± 3.7 kcal/mol (mean ± SD over 200 frames), followed by AURKB–taraxacin (−18.4 ± 2.7 kcal/mol) and KIF11–L-thyronine (−4.3 ± 4.9 kcal/mol). The favourable contribution in every complex came from gas-phase interactions (ΔG_gas_ = −61.6, −47.7 and −12.2 kcal/mol for CDK1, AURKB and KIF11), with van der Waals energy (−40.8, −28.9 and −8.4 kcal/mol) larger than electrostatic energy (−20.8, −18.8 and −3.9 kcal/mol), in line with the hydrophobic-dominated interaction patterns in Section 3.7. This was largely offset by unfavourable polar solvation (ΔE_GB_ = +35.9, +33.7 and +9.3 kcal/mol), while the non-polar solvation term contributed −5.3, −4.4 and −1.4 kcal/mol. The order CDK1 > AURKB > KIF11 is the same as the order of the docking scores (−10.7, −9.7 and −8.5 kcal/mol), but the spread between the complexes is much larger.

**Figure 14.**
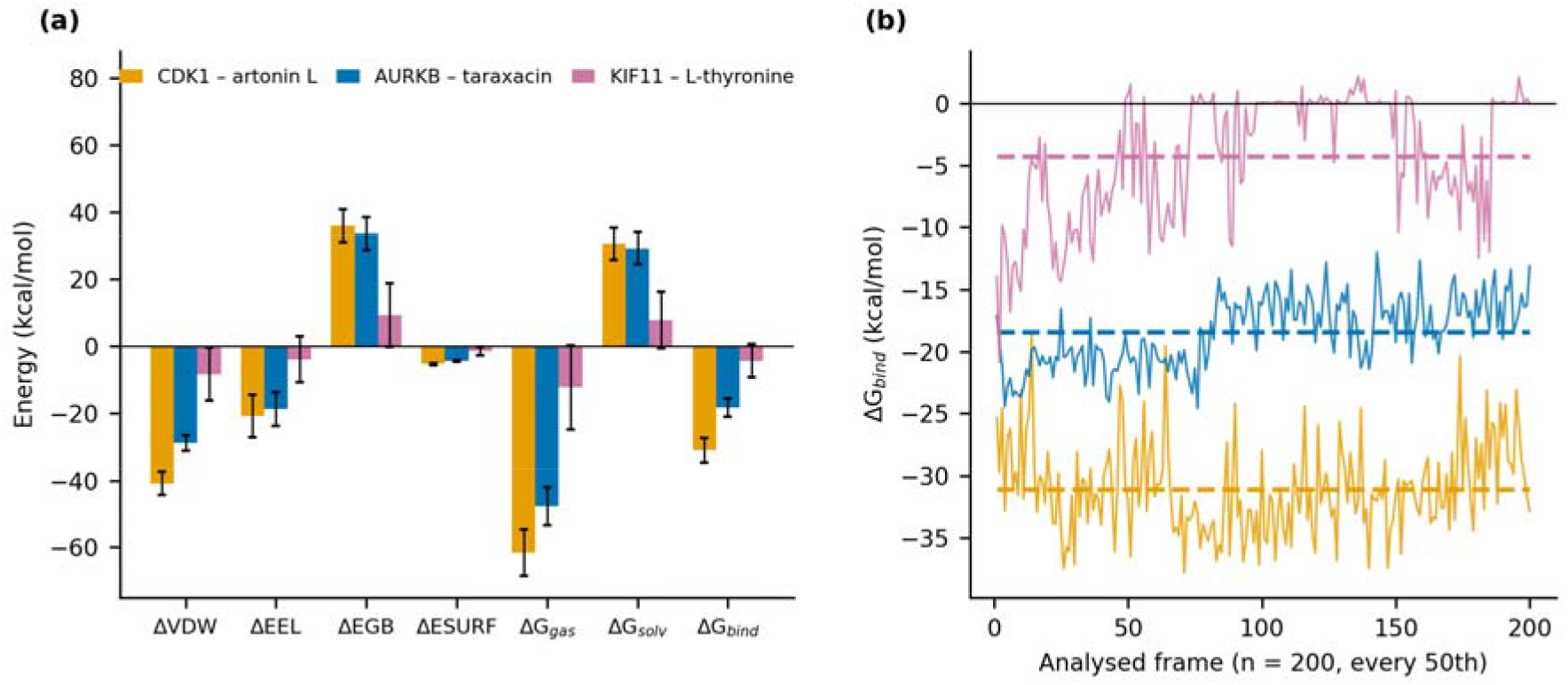
MM/GBSA binding free energies of the three protein–ligand complexes. (a) Energy components (mean ± SD over 200 frames). (b) Per-frame ΔG_bind_; dashed lines are the means. CDK1–artonin L (orange), AURKB–taraxacin (blue), KIF11–L-thyronine (pink). Entropy is not included.

**Table 8.** MM/GBSA binding free-energy components (kcal/mol; mean ± SD over 200 frames, single trajectory, entropy not included).

| **Component** | **CDK1–artonin L** | **AURKB–taraxacin** | **KIF11–L-thyronine** |
|---|---|---|---|
| **ΔE<sub>vdW</sub>** | −40.8 ± 3.5 | −28.9 ± 2.3 | −8.4 ± 7.9 |
| **ΔE<sub>ele</sub>** | −20.8 ± 6.3 | −18.8 ± 5.1 | −3.9 ± 6.8 |
| **ΔE<sub>GB</sub>** | 35.9 ± 4.8 | 33.7 ± 4.8 | 9.3 ± 9.5 |
| **ΔE<sub>SA</sub>** | −5.3 ± 0.4 | −4.4 ± 0.3 | −1.4 ± 1.3 |
| **ΔG<sub>gas</sub>** | −61.6 ± 6.9 | −47.7 ± 5.7 | −12.2 ± 12.5 |
| **ΔG<sub>solv</sub>** | 30.6 ± 4.7 | 29.3 ± 4.8 | 7.9 ± 8.4 |
| **ΔG<sub>bind</sub>** | −31.1 ± 3.7 | −18.4 ± 2.7 | −4.3 ± 4.9 |

The time courses differed in stability (Figure 15b). The CDK1 estimate was constant between the first and second halves of the analysed frames (−31.3 and −30.9 kcal/mol; lag-1 autocorrelation 0.25). The AURKB estimate weakened from −20.1 to −16.8 kcal/mol and the KIF11 estimate from −6.6 to −2.0 kcal/mol between the two halves. For KIF11, individual frames ranged from −20.9 to +2.2 kcal/mol, with 91 of the 200 frames above −2 kcal/mol and 60 above zero, which is consistent with weak and intermittent association of L-thyronine with the allosteric pocket (ligand–protein distances were not analysed here). Consecutive frames were correlated (lag-1 autocorrelation 0.67 for AURKB and 0.74 for KIF11), giving about 40 and 30 independent frames; the standard error of the mean accordingly rises from 0.19 to 0.44 kcal/mol for AURKB and from 0.35 to 0.89 kcal/mol for KIF11 (CDK1: 0.26 to 0.34 kcal/mol). Because entropy is not included and a single trajectory was analysed per complex, the absolute values tend to overestimate binding and should be used only to compare the complexes. The ΔG_bind_ of −4.3 kcal/mol for KIF11 is small relative to the uncertainty usually associated with MM/GBSA estimates and does not on its own support stable binding of L-thyronine, whereas the CDK1–artonin L value is both large and stable.

## 4. Discussion

This study set out to answer two questions the existing literature had left separate: whether the mitotic and cell-cycle hubs recurring across independent breast and ovarian cohorts represent a genuine shared vulnerability rather than an artifact read off two parallel gene lists, and whether any of them could be carried past a regulator list into structure-based drug screening. The hub-gene and GEPIA2 evidence answer the first question directly. CDK1, AURKB, CCNA2, and KIF11 were not only distilled from 396 DEGs shared between breast and ovarian cancer by PPI network topology, but independently confirmed as significantly upregulated in both BRCA and OV cohorts using GEPIA2, an expression-profiling platform built on TCGA tumor and GTEx normal-tissue data and maintained separately from the DEG/PPI pipeline that first flagged these genes. Together with TOP2A, this set of hubs converges functionally on G2/M-phase transition and mitotic spindle organization by GO/KEGG enrichment, consistent with CDK1’s established role as the G2/M checkpoint regulator and the Aurora kinases’ role in centrosome maturation and chromosome segregation. The second question, whether any hub can be carried into structure-based screening, was addressed through a sequential target- and ligand-prioritization strategy. Of the eight hub genes carried forward to docking, the three highest-ranked complexes, CDK1, AURKB and KIF11, were prioritized for the decoy-based validation and 100 ns molecular dynamics stages reported here. The ligand side of the panel underwent a parallel filtering process consistent with the study’s stated caution around unfamiliar chemical space: of 70,478 candidate phytochemicals drawn from FooDB, 233 survived sequential drug-likeness and structural-alert filtering, ML/QSAR-predicted bioactivity screening, and toxicity triage in Deep-PK, a deep-learning ADMET platform covering AMES mutagenicity, carcinogenicity, and drug-induced liver injury among its endpoints. Artonin L, taraxacin, and L-thyronine, the ligands retained in the three complexes discussed below, were therefore drawn from an already-filtered pool rather than selected post hoc from docking scores alone, though because IC50 prediction accuracy is known to degrade on chemically unfamiliar, food-derived scaffolds, this filtering step is treated here as a prioritization aid rather than a substitute for the binding-mode and dynamics evidence that follows.

The three prioritized complexes differed appreciably in how well their predicted binding modes could be reconciled with independent literature. For CDK1, the ATP-competitive contacts recovered for CID 44258662 (artonin L), spanning Phe80, Ile10, Val18 and Leu135, match essentially residue-for-residue the canonical CDK1 ATP-binding cleft defined in independent pharmacophore and virtual-screening studies,^113,114^ and the artonin scaffold itself has repeatedly been shown to induce apoptosis together with S-phase cell-cycle arrest in cancer cell lines, through mechanisms compatible with, if not specific to, cell-cycle kinase engagement.^115,116^ A comparable convergence was seen for AURKB: the hydrogen bond to Phe219 and the surrounding hydrophobic network (Phe88, Val91, Leu207) recovered for CID 5241825 (taraxacin) reproduces residues independently identified as central to Aurora B inhibitor binding in the same QSAR/molecular-dynamics study,^117^ lending plausibility to the pose despite the absence of any direct pharmacological data on taraxacin itself; more broadly, guaianolide sesquiterpene lactones are a well-documented class of natural cytotoxic agents.^118^ The KIF11 complex with CID 5461103 departed from this hydrophobic-dominant pattern, instead anchored through Glu116 and Leu214, both established anchor residues of the pharmacologically validated α2/loop L5/α3 allosteric pocket engaged by clinical-stage Eg5 inhibitors including ispinesib and S-trityl-L-cysteine.^119,120^ This correspondence is notable given that the ligand itself, L-thyronine, is the non-iodinated parent scaffold of the thyroid hormones rather than a characterized bioactive hormone in its own right, and its behavior cannot be assumed to mirror that of thyroxine (T4), whose tumor-relevant, non-genomic action at the integrin αvβ3 receptor is documented specifically for the iodinated hormone.^121^

The molecular dynamics data were broadly consistent with this picture. CDK1 and KIF11 each showed rapid RMSD equilibration to tight plateaus, restricted PCA phase space, and densely coupled DCCM networks, consistent with well-anchored ligands occupying validated pockets, even though KIF11 showed weaker decoy discrimination and higher solvent exposure than CDK1, the latter plausibly reflecting the intrinsically more open character of part of the Eg5 allosteric site rather than a poor pose. AURKB combined the largest RMSD/RMSF excursions in the panel with the lowest radius of gyration, lowest SASA and a PCA signature of transitions between discrete sub-clusters rather than progressive unfolding, a combination more consistent with conformational selection among related compact states than with genuine destabilization. One plausible contributor is that nearly all resolved Aurora B structures, including the template used in independent AURKB docking and QSAR studies are of the kinase in complex with its activating partner INCENP as part of the chromosomal passenger complex, rather than the isolated kinase domain;^122^ the present simulation used AURKB alone, additional flexibility relative to the INCENP-bound state would be a plausible, checkable consequence of the missing partner rather than evidence against the docking pose itself.

The additional validation separates the three complexes more sharply than the docking scores alone. The reported scores and binding modes are reproducible: re-docking returned all three scores to within 0.7 kcal/mol, every reported contact residue reappeared in the re-docked poses, and all of those residues line the crystallographic ligand pocket of their target, so the poses obtained without specifying a site occupy the experimentally characterised binding sites. What the benchmark constrains is the meaning of the scores rather than the poses. Against measured ChEMBL actives and inactives the Vina score was only weakly discriminative (ROC-AUC 0.57–0.72) and, at CDK1 and AURKB, no better than heavy-atom count alone. On that scale artonin L at CDK1 is the strongest candidate: it out-scored 80% of known CDK1 actives, its pose is recovered under every search condition tested, and its MM/GBSA free energy is the largest and the only one without drift between the two halves of the analysed frames (−31.1 kcal/mol). Taraxacin at AURKB is supported by a reproducible score and pose in the ATP cleft and by a mid-range standing among measured AURKB actives that improves to the top of the range once ligand size is taken into account, but its MM/GBSA energy weakened over the analysed frames (−20.1 to −16.8 kcal/mol), so the complex is plausible but less firmly established. L-Thyronine at KIF11 is the least well supported. Its pose falls in the loop-L5 allosteric pocket of Eg5 and its benchmark standing matches that of taraxacin, but its MM/GBSA free energy was small and unstable (−4.3 kcal/mol, weakening from −6.6 to −2.0 kcal/mol between the two halves of the analysed frames, with 60 of the 200 frames above zero), which does not support persistent binding. The backbone stability reported for the KIF11 and AURKB simulations describes the protein rather than the ligand and should not be read as evidence of stable ligand binding.

Taken together, CDK1 is the most robustly supported complex in this panel on docking score and conformational stability, while AURKB shows the strongest discrimination against property-matched decoys (Z = −1.77 and the 98th percentile, versus Z = −1.50 and the 94th percentile for CDK1) and is comparably well supported at the level of pocket recognition, with its greater conformational flexibility attributable to a plausible, checkable structural cause rather than to a poorly defined pose. KIF11 rests on a specific, literature-validated allosteric binding mode, but its decoy discrimination was the weakest of the three credible complexes, and the ligand itself has no independent history as a bioactive compound, so this result is best presented as hypothesis-generating rather than confirmatory.

## 5. Conclusion

This study pooled six independent breast and ovarian carcinoma cohorts into a single analytical pipeline and recovered 396 shared differentially expressed genes, from which network topology prioritized ten hub genes spanning the mitotic, apoptotic and PI3K–AKT–FOXO programs. Seven of the ten reproduced in independent TCGA/GTEx data, and functional enrichment placed the signature on G2/M transition, mitotic spindle organization and p53 signalling, with TP53, STAT3, TFAP2A, FOXC1 and GATA2, together with the miR-15/16, miR-21, miR-26a and let-7a families, as principal upstream regulators. None of these genes is new to either disease individually. What the pipeline adds is that their shared status is derived from one integrated comparison rather than inferred across separate single-disease reports, and that the surviving hubs were carried forward into structure-based screening. Of 70,478 food-derived compounds, 233 survived drug-likeness, bioactivity and toxicity filtering and were docked against eight druggable hubs. CDK1 emerged as the best-supported target: artonin L occupied the canonical ATP cleft, discriminated cleanly from property-matched decoys, and held a stable trajectory across the simulation. AURKB was comparably well recognized at the pocket level, with its greater conformational mobility attributable to simulation of the kinase in the absence of its INCENP partner. KIF11 bound at the pharmacologically validated α2/L5/α3 allosteric site but showed the weakest decoy separation, with a ligand of no established bioactivity. These findings are computational and hypothesis-generating. They nominate CDK1 as the priority candidate for in vitro kinase-inhibition and cell-viability testing in paired breast and ovarian models, and provide a reusable template for extending cross-cancer transcriptomic prioritization into natural-product structure-based discovery.

## Limitations

Several limitations qualify these findings. First, the study is entirely computational: no wet-laboratory validation of target expression, compound binding or cellular activity was undertaken, so the prioritized complexes remain hypotheses rather than demonstrated interactions

## Abbreviations

ADMET: absorption, distribution, metabolism, excretion and toxicity
BRCA: breast invasive carcinoma
cDEG: common differentially expressed gene
DCCM: dynamic cross-correlation matrix
DEG: differentially expressed gene
DILI: drug-induced liver injury
GEO: Gene Expression Omnibus
GO: Gene Ontology
GRN: gene regulatory network
GTEx: Genotype-Tissue Expression project
KEGG: Kyoto Encyclopedia of Genes and Genomes
LUDe: LIDeB Useful Decoys
MD: molecular dynamics
MM/GBSA: molecular mechanics/generalised Born surface area
NPT: isothermal–isobaric ensemble
NVT: canonical ensemble
OV: ovarian serous cystadenocarcinoma
PAINS: pan-assay interference compounds
PCA: principal component analysis
PPI: protein–protein interaction
QSAR: quantitative structure–activity relationship
Rg: radius of gyration
RMSD: root-mean-square deviation
RMSF: root-mean-square fluctuation
ROC-AUC: area under the receiver operating characteristic curve
SASA: solvent-accessible surface area
sDEG: shared differentially expressed gene
sKG: shared key gene
TCGA: The Cancer Genome Atlas
TF: transcription factor

## Author contributions

Conceptualization: MJT, AS. Data curation: MJT. Formal analysis: MJT, MIHA. Investigation: MJT. Methodology: MJT. Software: MJT, CB. Visualization: MJT, MIHA, AS. Writing – original draft: MJT, MIHA. Writing – review and editing: MJT, AS. All authors read and approved the final manuscript.

## Data availability

All data analysed in this study are publicly available. The gene expression datasets were obtained from the NCBI Gene Expression Omnibus under accessions GSE45827, GSE65194, GSE14407, GSE18520, GSE29450 and GSE54388 (https://www.ncbi.nlm.nih.gov/geo/). Validation expression data were accessed through GEPIA2 (http://gepia2.cancer-pku.cn/), which draws on TCGA and GTEx. Protein–protein interaction data were retrieved from STRING (https://string-db.org/); transcription-factor and miRNA interactions from ChEA3 and TarBase v9.0 via NetworkAnalyst 3.0 (https://www.networkanalyst.ca/). The phytochemical library was downloaded from FooDB (https://foodb.ca/) and bioactivity training data from ChEMBL release 37 (https://www.ebi.ac.uk/chembl/). Receptor structures were obtained from the RCSB Protein Data Bank under accessions 1MQ4, 4AF3, 1XOX, 4BCN, 4Y72, 8A62, 3K5E and 9BQ6. Supplementary data are provided with this article.

## Ethics approval and consent to participate

Not applicable. This study is a secondary computational analysis of publicly available, de-identified datasets and did not involve new human participants, human material or animal experimentation.

## Declaration of generative AI and AI-assisted technologies in the manuscript preparation process

During the preparation of this work, the authors used AI-assisted tools for language polishing, grammatical correction, and improving the clarity and readability of the manuscript.

