## Supplementary Table 1 for "Convergence of Independently Established Oncogenes in a Combined Breast and Ovarian Cancer Model: Hub Gene Validation and identification of potential Food-Derived therapeutics"

| Supplementary Table 1 Top enriched Gene Ontology (GO) terms and KEGG pathways for the ten prioritised hub genes. | | | | |
| --- | --- | --- | --- | --- |
| **Categories** | **GO ID / Overlap** | **GO-Terms / Pathway** | **Adj. P-value** | **Associated hub genes** |
| Biological Process (BPs) | GO:0044839 | Cell Cycle G2/M Phase Transition | 7.02E-09 | CCNA2; CDK1; BIRC5; AURKB; AURKA |
|  | GO:1902850 | Microtubule Cytoskeleton Organization Involved in Mitosis | 1.40E-08 | CDK1; BIRC5; KIF11; AURKB; AURKA |
|  | GO:0007052 | Mitotic Spindle Organization | 8.05E-06 | BIRC5; KIF11; AURKB; AURKA |
|  | GO:0042981 | Regulation of Apoptotic Process | 3.03E-04 | TOP2A; BCL2; CDK1; BIRC5; FOXO1 |
|  | GO:0006974 | DNA Damage Response | 6.60E-04 | TOP2A; BCL2; CDK1; FOXO1 |
| Molecular Function (MFs) | GO:0046982 | Protein Heterodimerization Activity | 1.25E-04 | TOP2A; BCL2; BIRC5; AURKA |
|  | GO:0019900 | Kinase Binding | 1.18E-03 | CCNA2; KIF11; AURKB; AURKA |
|  | GO:0004712 | Protein Serine/Threonine/Tyrosine Kinase Activity | 1.18E-03 | AURKB; AURKA |
|  | GO:0019901 | Protein Kinase Binding | 1.18E-03 | TOP2A; CCNA2; KIF11; AURKA |
|  | GO:0004674 | Protein Serine/Threonine Kinase Activity | 7.75E-03 | CDK1; AURKB; AURKA |
| Cellular Components (CCs) | GO:0005876 | Spindle Microtubule | 5.89E-09 | CDK1; BIRC5; KIF11; AURKB; AURKA |
|  | GO:0005874 | Microtubule | 2.71E-07 | CDK1; BIRC5; KIF11; AURKB; AURKA |
|  | GO:0005634 | Nucleus | 1.44E-05 | TOP2A; CCNA2; BCL2; CDK1; PTEN; BIRC5; KIF11; AURKB; FOXO1; AURKA |
|  | GO:0015630 | Microtubule Cytoskeleton | 9.82E-05 | BIRC5; KIF11; AURKB; AURKA |
|  | GO:0000307 | Cyclin-dependent Protein Kinase Holoenzyme Complex | 8.42E-04 | CCNA2; CDK1 |
| KEGG Pathway | 4/156 | Cellular Senescence | 5.41E-05 | CCNA2; CDK1; PTEN; FOXO1 |
|  | 5/525 | Pathways in Cancer | 1.04E-04 | CCNA2; BCL2; PTEN; BIRC5; FOXO1 |
|  | 3/75 | p53 Signaling Pathway | 1.49E-04 | BCL2; CDK1; PTEN |
|  | 3/103 | Prostate Cancer | 2.68E-04 | BCL2; PTEN; FOXO1 |
|  | 3/157 | Cell Cycle | 6.32E-04 | CCNA2; CDK1; AURKB |
